# Behavioral Time Scale Synaptic Plasticity (BTSP) endows binding of distributed representations with flexible retrieval options

**DOI:** 10.1101/2025.05.15.654220

**Authors:** Chengting Yu, Yukun Yang, Yujie Wu, Aili Wang, Wolfgang Maass

**Affiliations:** Institute of Machine Learning and Neural Computation, Graz University of Technology, Graz, Austria; College of Information Science and Electronic Engineering, Zhejiang University, Hangzhou, China; Department of Computing, The Hong Kong Polytechnic University, Hong Kong SAR, China

**Author notes:** Email addresses.

## Abstract

Human reasoning depends on reusing pieces of information by binding them together in new ways, thereby “making infinite uses of finite means” (Alexander von Humboldt). Needed for that is a binding mechanism that enables fast composition and decomposition of tokens of information. Binding can easily be implemented in symbolic computations through parentheses and ordering of symbols. But it is a highly nontrivial operation for distributed representations, where the tokens are encoded by activity patterns in large neural networks or large language models, or more abstractly, by a high dimensional vector. Vector Symbolic Architectures (VSAs) provide partial solutions, but are lacking the flexibility of the brain in information retrieval, e.g. retrieving the tokens from a composed representation or retrieval of a composed representation by just providing some tokens as a cue. We show that a mechanism which the brain employs for binding distributed representations, Behavioral Time Scale Synaptic Plasticity (BTSP), overcomes these deficiencies. In particular, it combines binding with attractor features that make information retrieval substantially more flexible and robust. We evaluate its performance on various applications, including encoding and decoding complex visual scenes and hierarchical binding. We also show that it enhances models for natural language processing and abstract brain computation. BTSP-based binding only requires binary synaptic weights and simple local synaptic plasticity, and can therefore easily be implemented through in-memory computing or other innovative designs for energy-efficient AI implementations.

## 1 Introduction

Most higher cognitive functions of the human brain rely on some form of compositional or symbolic computation, where entities such as symbols, objects, persons, events, as well as features of them and relations between them, are bound together into composed representations. One commonly refers to these entities as components, tokens, words, or items. Like a sentence that is composed of words in natural language, these composed representations acquire new meanings and functional roles that go beyond those of their individual components (Frankland and Greene, 2020a; Kurth-Nelson et al., 2023; Kazanina and Poeppel, 2023). Most neural network models and LLMs (large language models) have problems to reproduce this type of combinatorial coding and compositional computing, which has been argued to contribute to their deficits regarding transparency and trustworthiness (Campbell et al., 2024; Alonso et al., 2025).

An essential feature of these brain computations is that they are carried out on very sparse neural codes where the firing of just a few among millions of neurons encodes salient objects or concepts, and also structural information such as relations between them, temporal or spatial locations, or semantic roles of words in a sentence (Piantadosi et al., 2024). This perspective had inspired models for computation on high-dimensional (high-D) binary vectors that mostly consist of 0’s, such as the sparse distributed memory models of (Kanerva, 1988) and the holographic reduced representations of (Plate, 1995). These approaches inspired quite a bit of further work that gave rise to the research areas that are now commonly referred to as Hyperdimensional Computing (HDC) or Vector Symbolic Architectures (VSAs) (Kleyko et al., 2022, 2023). Both terms are viewed as synonyms, we use for simplicity only the term HDC. A prominent special case are the semantic pointer architecture that underlie the neural engineering framework of (Eliasmith and Anderson, 2003; Eliasmith et al., 2012). The challenge of binding distributed representations is discussed in (Kleyko et al., 2022). Not only the tokens, but also their grouping and order need to be captured by composed representations, similarly as brackets and symbolic order in symbolic representations.

The computational operations that have so far been carried out in HDC for binding high-D vectors, such as convolution or permutation, as well as component-wise addition, exclusive OR, or multiplication, are of an algebraic nature. These are hard to interpret as operations in neural networks of the brain. But more important are their functional deficits. Due to the mathematical fact that one cannot recover the component bits from a sum, product, or exclusive OR of two bits, decoding of the tokens from a composed representation is in general impossible in HDC, unless some of the components are given as cue. Only one method has been proposed in the HDC framework which achieves something of this type: resonator networks (Frady et al., 2020; Kent et al., 2020; Renner et al., 2024). But these resonator networks need to be given a codebook of all possible tokens that might occur in a composed representation. They then go through an iterative process through all possible combinations of tokens that are consistent with a given composed representation. Hence they are less suitable if the number of possible tokens is very large, which occurs in real world applications, and also for hierarchical binding. They are also less suitable if fast recovery of tokens is needed because of the duration of this iterative process. An additional deficit of previous considered schemes for binding in HDC is the incapability to reproduce the composed representation or other tokens in its by presenting just one or a few of them, without providing also the composed representation.

We present here a new approach to HDC that overcomes these deficits. Algebraic operations are replaced here by a mechanism that the brain employs for binding, Behavioral Time Scale Synaptic Plasticity (BTSP) (Bittner et al., 2015, 2017; Zhao et al., 2022). BTSP was more recently discovered than Hebbian plasticity or STDP (Spike Timing Dependent Plasticity). Furthermore, it is one of very few synaptic plasticity mechanisms that could be demonstrated in-vivo, even in the awake and operating adult brain, see (Magee and Grienberger, 2020; Chéreau et al., 2022) for reviews. In particular, experimental data show that the brain uses BTSP for binding information (Bittner et al., 2015; Zhao et al., 2022).

In contrast to other synaptic plasticity mechanisms, BTSP is a one-shot mechanism that modifies synaptic weights instantly, after a single trial (Bittner et al., 2017). Another unique feature of BTSP is that it is triggered in area CA1 of the hippocampus by largely stochastic gating signals from another brain area, the entorhinal cortex (Grienberger and Magee, 2022). We exploit that this largely stochastic gating enables BTSP to combine functional benefits of random projections and random hashing, i.e., of widely used methods in computer science for distributing information over large memory systems, with additional features that are reminiscent of attractors in dynamical systems. More specifically, we show that this attractor feature of BTSP enables operations on composed representations in HDC, such us top-down unbinding and bottom-up unbinding without providing also the composed representation as cue, that brains can apparently accomplish, but which are not within reach of previously proposed HDC methods for binding. Furthermore, we show that these functional contributions of BTSP can be captured with a simple rule for BTSP that only requires binary weights (Wu and Maass, 2025). This enables implementation of the BTSP-enhanced version of HDC by in-memory computing with arrays of highly energy-efficient memristors even if these can assume only a few different conductance states in a reliable manner. Important examples for that are phase-change memristors as described in (Wang et al., 2020; Khaddam-Aljameh et al., 2022).

The BTSP-enhanced version of HDC also contributes new mechanisms and functional capabilities to models that aim at reproducing higher cognitive functions of the brain in simplified computational models that are analytically tractable, see e.g. (Valiant, 2000; Papadimitriou et al., 2020). Experimental data suggest that these tokens of meaning are encoded by very sparse distributed assemblies of neurons that fire when a particular concept is invoked. A prominent example are assemblies of concept cells (Quiroga, 2012; Franch et al., 2025). Hence, if the current firing activity of all neurons in the brain is represented by a high-D vector, the activation of all neurons in an assembly is represented by a sparse high-D vector. A fundamental operation in these models is the binding of two assemblies, or of an assembly of concept cells with a assembly of neurons that encodes structural information (Müller et al., 2020; Kazanina and Poeppel, 2023). Hence binding of high-D vectors is a key issue in these abstract models for brain computation (Piantadosi et al., 2024). This binding operation has commonly been modeled by an iterative process, where reciprocal connections between neurons of two assemblies are modified by STDP (Valiant, 2000; Papadimitriou et al., 2020; Papadimitriou and Friederici, 2022). But such an iterative process is neither consistent with the capability of brains to form composed representations instantly, nor with the large number of iterations that are needed to modify in experimental studies a synaptic weight by STDP (Froemke et al., 2010). Our BTSP-enhanced version of HDC provides a simpler, biologically more plausible, and functionally advantageous binding operation for these models. Interestingly, this binding operation does not modify synaptic weights between the neurons of two assemblies in the neocortex, but synaptic weights to and from other large pools of neurons, such as the hippocampus. In other words, the BTSP-enhanced variant of HDC supports a model for brain computation that is consistent with the “indexing theory”, whereby hippocampal neurons form conjunctive codes that act as pointers to item representations in the neocortex. Recent experimental work has shown that this indexing theory is consistent with the way how the human brain carries out binding operations (Kolibius et al., 2023).

BTSP-enhanced HDC also provides a fresh perspective for computational models of natural language processing in the human brain. For example, it had been proposed that the grammar of natural languages only requires a single operation, the Merge operation (Chomsky et al., 2023; Liu et al., 2023). So far the Merge operation has been modeled as iterative process via STDP (Papadimitriou and Friederici, 2022). But this appears to be in conflict with the need to carry out Merge instantaneously during language processing. Hence BTSP-enhanced HDC appears to provide a more suitable framework for modeling the implementation of the Merge operation in neural networks of the brain.

We will first review fundamental properties of experimental data and models for BTSP that are essential for its binding capability, in particular the attractor features that it contributes. We then demonstrate the capability of BTSP to enhance fundamental operations of compositional computing, such as top-down unbinding and bottom-up unbinding. Subsequently we compare the unbinding performance for BTSP-based binding with that of HDC-based binding. We then show for BTSP-based binding that bottom-up unbinding is also possible if given cues are ambiguous, i.e., may have several possible solutions. Finally, we examine the capability of BTSP to support hierarchical binding and unbinding. In the last section we discuss possible applications of BTSP-based binding in an assembly-based brain calculus for higher cognitive functions, in particular for language processing.

## 2 Results

### 2.1 Fundamental properties of BTSP that are essential for its binding capability

Experimental data show that the brain uses BTSP to form conjunctive representations of diverse content that is encoded by sparse activity in areas CA3 (“input neurons”) through sparse representations in area CA1 (“memory neurons”) (Bittner et al., 2015, 2017; Zhao et al., 2022), see Fig. 1A for a network scheme. We use here the term “memory neurons” for those neurons in area CA1 that are recruited for composed representations in order maintain consistency with the notation in (Wu and Maass, 2025). BTSP differs in several fundamental properties from plasticity rules that are commonly considered in neural network models. One is, that it does not require repeated representations of the same input or input/output pairing as STDP or other commonly considered plasticity rules. Rather, it creates new representations in a single shot. Obviously, this property is essential for binding and compositional computing in general, since new combinations of familiar items, referred to as words in the following, have to be learned in one shot, for example when we hear or read a sentence.

**Fig. 1:**
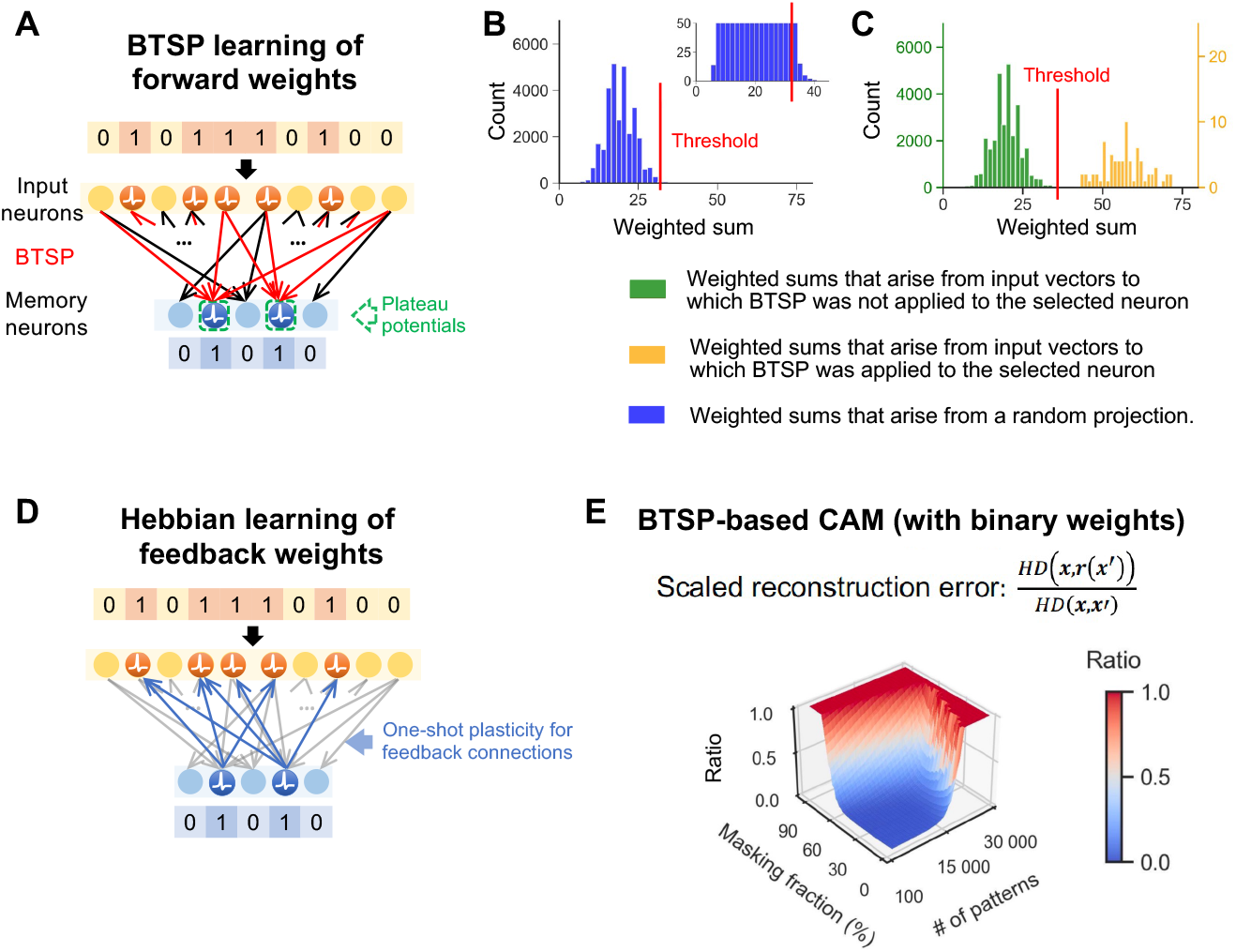
Fundamental properties of BTSP. **(A)** BTSP transforms an input string of bits, represented by firing or non-firing of input neurons, into a corresponding representation of an output bit string by memory neurons through one-shot learning, with synaptic plasticity restricted to memory neurons that receive a gating signal (plateau potential). **(B)** Distribution of weighted sums at a generic memory neuron that results from applying a fixed random projection (a fixed stochastic binary matrix, whose entries are independent of the input vectors) to 30k input vectors. No matter how one sets the firing threshold of the neuron (see insert for a blow-up), numerous input vectors cause weighted sums that are close to the threshold, and therefore may cross the threshold when the input vector is slightly perturbed. **(C)** If the binary synaptic weights result instead from applications of BTSP to the same list of input patterns, the distribution of weighted sums becomes bi-modal: One mode (green) arises from input patterns for which BTSP was not applied to this neuron, and another mode (yellow) from input patterns for which BTSP was applied to this neuron. This bi-modal distribution enables a choice of the firing threshold that is robust to small changes in the input pattern. This noise-robustness lies at the heart of the attractor features of composed representations in BTSPbased HDC. Panels B and C are reprinted from (Wu and Maass, 2025). **(D)** In order to be able to reconstruct an input vector from a composed representation, also synaptic weights of connections from the memory neurons to the input neurons have to be assigned. One-shot learning with Hebbian plasticity suffices, due to the sparsity of both bit strings. Note that Hebbian plasticity, rather than BTSP, is suitable for this type of synaptic plasticity because the values in the input pattern can be seen as postsynaptic activity in this case of self-supervised learning. **(E)** The noise robustness of BTSP from panel C endows composed representations with attractor features that, in conjunction with the feedback connections from panel D, create a content addressable memory (CAM) for those input patterns for which BTSP was applied. This panel was copied from (Wu and Maass, 2025).

Another fundamental difference between BTSP and previously considered learning rules such as Hebb or STDP is that it does not depend on postsynaptic neuronal activity. A positive influence of postsynaptic firing on LTP would entail that the same neurons are recruited for many composed representations, since their recruitment for some of them would increase their incoming weights through LTP, and hence increase the chance that they also fire during the formation of further composed representations. Instead, BTSP is gated by dendritic plateau potentials in the postsynaptic neuron that are triggered by largely stochastic synaptic inputs from layer 3 of the entorhinal cortex (Grienberger and Magee, 2022). The stochastic feature of these gating signal induces a more uniform distribution of the recruitment of memory neurons for composed representations. These gating signals open the gate for synaptic plasticity through BTSP for several seconds before and after its onset(Bittner et al., 2017; Magee and Grienberger, 2020; Milstein et al., 2021). We need this long plasticity in our binding model only implicitly, since we work in our subsequent evaluations for simplicity with batch inputs *x* whose bits arrive simultaneously in the model. In the brain the long plasticity window of BTSP allows to collect content information that arrives from several brain areas on a behavioral time scale of seconds, and this long integration time for plasticity would also be useful for applications in artificial devices in order to bind input components from different sensory modalities and other network modules that do not arrive simultaneously, or for binding a sequence of frames together as suggested by recent experimental from the human hippocampus (John et al., 2025).

It was shown in (Wu and Maass, 2025) that the following simple plasticity rule with binary weights *w*_*i*_ captures key features of BTSP in the brain. Synaptic plasticity takes place only when its plasticity window is opened by a stochastic gating signal, and if there is in addition presynaptic activity, i.e., *x*_*i*_ = 1, where *x*_*i*_ denotes presynaptic activity (encoded by a binary state: *active* = 1, *inactive* = 0). The new value of the weight is then

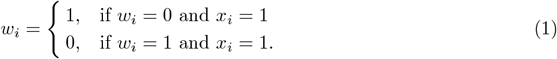

According to this rule, synaptic weights undergo Long-Term Potentiation (LTP) or Long-Term Depression (LTD), in dependence of their preceding value, and of the presynaptic activity. The weight-dependent aspect of BTSP, that had been elucidated in (Milstein et al., 2021), may look surprising from the functional perspective, since it could in principle weaken the capacity of the network. But if brain-like sparse input patterns and gating signals are used for BTSP, most weights have value 0 even after a large numbers of patterns have been stored Wu and Maass (2025). Consequently, LTD is applied only rarely. Nevertheless, it was shown in (Wu and Maass, 2025) that this rare application of LTD is helpful for maintaining a low fraction of non-zero synaptic weights, and for reproducing the repulsion effect of the human brain.

Whereas BTSP looks from this perspective more similar to random hashing, it combines the benefits of random hashing with a key property of Hebbian learning that random hashing cannot provide: It creates attractor properties for stored memories, so that these can also be recalled with partial or perturbed versions of the original content. An explanation on the mechanistic level is provided by Fig. 1 B and C. Its functional benefits become obvious if one trains simultaneously with the feedforward connections through BTSP also backwards connections from memory neurons to input neurons (Fig. 1D). These backwards connections can be viewed as parsimonious model for the well-known big-loop recurrence of the brain between CA1 and CA3 (Koster et al., 2018). Since sparse gating signals for BTSP induce sparse representations on the memory layer, and since we work with sparse input patterns, it suffices to use just binary weights also for the feedback connections, and to train them with a simple plasticity rule with flips the weight value if both the pre- and postsynaptic neuron are firing (i.e., assume value 1). This plasticity rule is closely related to a variant of BTSP in area CA3 Li et al. (2024), which also relies on postsynaptic firing, and can therefore be viewed as a variant of Hebbian learning. If one combines the feedforward synaptic plasticity of Fig. 1A with the plasticity of feedback connections of Fig. 1D one arrives at a functionally powerful model for content addressable memory (CAM), see Fig. 1E: The Hamming distance (HD) between a pattern *x* that had been learnt through BTSP, and a partially masked variant *x*^*′*^ of this pattern is reduced when 4 *x*^*′*^ is replaced by the reconstructed version *r*(*x*^*′*^) of *x*^*′*^ that results from applying the feedforward weights from Fig. 1A in conjunction with the feedback weights from Fig. 1D. The reconstruction *r*(*x*^*′*^) arises already if forward and feedback weights are applied just once, i.e., no further recurrent network dynamics has to be examined.

Fig. 1E, which is copied from (Wu and Maass, 2025), shows that a fairly large masking fraction of 50% of the bits of *x* can be tolerated in variants *x*^*′*^ of *x*, in the sense that most of the missing bits are correctly reconstructed by *r*(*x*^*′*^). This content-addressable-memory (CAM) property that is induced by BTSP will be useful for improving the fidelity of top-down unbinding that we consider in the next section.

We will use in the subsequent evaluations of properties of BTSP-based binding largely the same brain-based values of hyperparameters as in Wu and Maass (2025). Input patterns are sparse binary vectors with a fraction *f*_*p*_ = 0.005 of 1’s. This matches the estimate of the sparsity of input patterns that CA1 receives from area CA3 according to the experimental data from Guzman et al. (2016). A memory neuron receives a gating signal with probability *f*_*q*_ = 0.0025 for a given input pattern, as suggested by the experimental data of (Grienberger and Magee, 2022). To be precise, we use here for simplicity the deterministic version (“core BTSP”) of (Wu and Maass, 2025), which is according to Fig. S1 of that paper for this value of *f*_*q*_ functionally equivalent to the stochastic version with *f*_*q*_ = 0.005. Note that *f*_*q*_ largely determines the sparsity of composed representations in the memory layer. Results of control experiments for different values of *f*_*p*_ and *f*_*q*_ can be found in Fig. S1 of the Supplement.

### 2.2 Binding

There exist various notions of binding in the literature. We will use binding in the sense of standard literature in cognitive science, neuroscience, and computational science. Binding of words into a sentence in natural language, or of different visual objects together with their features and positions in a complex visual scene, are the most common examples in neuroscience and cognitive science. The review (Frankland and Greene, 2020a) discusses the binding problem also more generally in terms of Fodor’s “language of thought hypothesis”, according to which our minds employ an amodal, language-like system for combining and recombining simple concepts to form more complex thoughts. The more recent review (Kurth-Nelson et al., 2023) also discusses compositional computations in the brain where entities are assembled into relationally bound neural codes to derive qualitatively new knowledge: Each entity is transiently bound to a representation of its role in the compound, which specifies how the element functions as part of the whole. In Fig. 1C, D they allude to binding of words to their role in a sentence, and binding of these compound representations into stories as a higher-level composition. Similar desirable functional properties of binding are discussed from the perspective of computational sciences in (Kleyko et al., 2022) under “Binding to address challenges of conventional connectionists representations”.

The definition of binding that we use agrees with these general definitions. We define binding as a function that maps a given list of high-D binary vectors (codes for the tokens) onto other high-D binary vectors, the composed representations. In BTSP-based binding the high-D input vectors A, B, … that encode components are concatenated and presented as binary activation pattern in an array of input neurons, see Fig. 2A. The output of the binding function, the composed representation, is defined by the resulting activity of binary neurons in the memory layer after application of BTSP, see Fig. 2A, C. The possibility to (approximately) invert this function, which we call top-down unbinding, is provided by a simultaneous application of a 1-shot Hebbian learning rule to the synaptic connections from the memory neurons to the input neurons (Fig. 2B). A closer look shows that this binding and unbinding operation depends on the order in which different binding demands arise. But our results show that this order is not salient for binding and unbinding performance if one applies it to brain-like sparse high-D vectors.

**Fig. 2:**
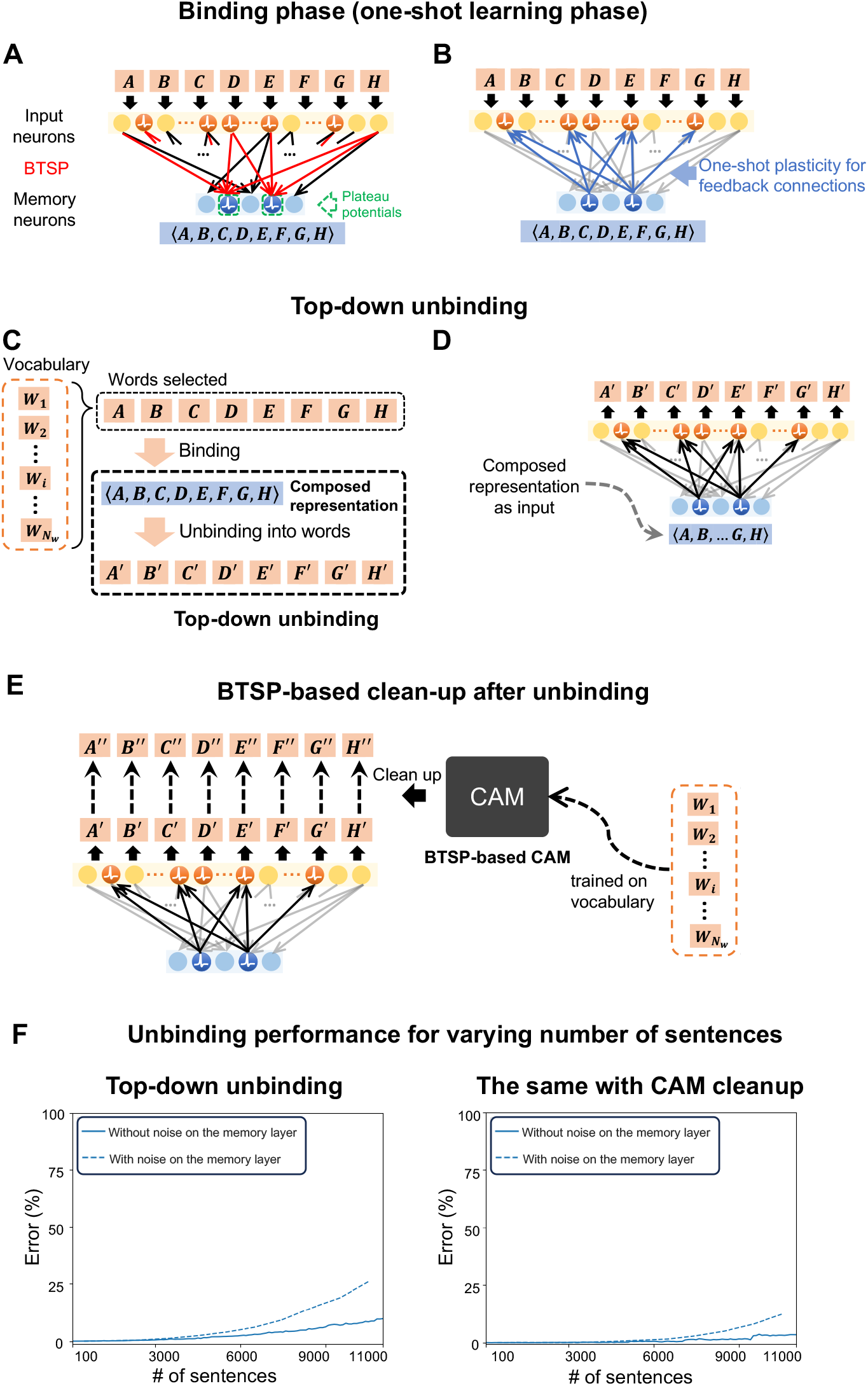
Top-down unbinding from a composed representation created through BTSP. We considered here the case where composed representations were created through BTSP from sequences of 8 words (each represented by sparse binary vectors). BTSP was applied sequentially to a sequence of *K* different combinations of 8 words, where K, the considered number of “sentences”, varied between 100 and 11,000. **(A**,**B)** For each of these sentences, BTSP was applied to the forward connections, and a Hebbian rule simultaneously to feedback connections, according to the general scheme of Fig. 1 A and D. **(C**,**D)** General scheme for binding 8 words into a composed representation, and for decoding these words from the composed representation. **(E)** Scheme for cleaning up decoded words A’, B’, ..by applying a BTSP-based CAM that was trained for the chosen vocabulary of 1000 words. **(F)** The resulting error is shown both without and with a cleanup of word codes via a BTSP-based CAM, as indicated in panel E. Decoding performance is good for up to 3000 sentences without a clean-up, and for about 7000 with the CAM. Furthermore, the dotted lines show that noise that flips the output of a memory neuron with probability 0.0025 affects the performance only slightly, in spite of the fact that this level of noise is on the same scale as the density of 1’s in composed representations.

### 2.3 Top-down unbinding

We apply in the following BTSP binding to content items (words) that are encoded by sparse binary vectors of length 6000. In view of the chosen brain-like sparsity level of *f*_*p*_ = 0.005 this amounts to an average number of 30 bits with value 1 in a word. We focus on binding processes where 8 words are bound together into a composed representation. Each composed representations (represented by neurons in the memory layer that fire) is a bit vectors of length 12,000, see the network scheme in Fig. 2A.

A key property of any type of binding scheme is to what extent items that are bound together can be recovered from a composed representation (top-down unbinding). Strangely enough, none of the numerous schemes for HDC enables that; one always needs to provide in addition to the composed representation a substantial fraction of its content items in order to recover the other ones, see the reviews Kleyko et al. (2022, 2023). In contrast, for binding via BTSP the simultaneously trained feedback connections (see Fig. 2B) enable top-down unbinding without providing any content items, see Fig. 2 C, D. We considered here the case where 8 words where bound into a composed representation in one shot through BTSP, for various numbers of sentences between 100 and 11,000.

The recovered noisy versions A’, B’, … of the words A, B, … have in general minor errors, see the left plot of Fig. 2F. But this error can be substantially reduced according to the right plot in Fig. 2F if one applies to each recovered word A’, B’,… a BTSP-based CAM that was trained for the underlying vocabulary of 1000 words. The plot shows that the resulting cleaned-up words A”, B”, .. have even for large numbers of sentences very little errors in comparison with the original words. We used in Fig. 2F the quotient of all incorrect bits in the decoded 8 words divided by the average number of 1’s in a sentence (given in %). Note that just counting the fraction of incorrect bits among all decoded bits would provide a too optimistic measurement: A decoding result consisting of just 0^*′*^*s* would already fair well, due to the fact that the vectors are very sparse (99.5% of them have value 0).

If one uses symbolic instead of distributed representations, there exists a trivial method for binding content vectors (words) into composed representations (sentences) that humans use in written language: One simply appends the codes for words (with blank spaces as separators). This method perfectly supports top-down decomposition, but it does not support bottom-up decomposition, discussed in the next section, because it has no attractor feature. In addition, this trivial method induces a sometimes undesirable growth of the length of coding vectors when one iterates the binding process. This can be avoided with BTSP-binding (see section 2.5).

### 2.4 Bottom-up unbinding

We consider here the same process for forming composed representations via BTSP as in the preceding section. But we examine now to what extent their content can be recovered without giving the composed representation as cue. Note that current HDC approaches support unbinding of composed representations only if both the composed representation and at least 1/2 of the content items are provided as cues. The latter is needed because in order to recover an input vector from a componentwise sum, product, or exclusive OR of two input vectors, the other input vector has to be provided as cue (in addition to the composed representation). Surprisingly, even without providing the composed representation as cue, less than 1/2 of the content items suffice as cue. For example, Fig. 3 A - C consider the case where just 2 out of 8 words are provided as cues. Fig. 3A illustrates a generic example, where just the words B and D are provided as cues. The same unbinding process as in Fig. 2 C, D generates from the resulting activity vector M(B, D) on the memory layer an approximation A’, B’, C’, D’, E’, F’, G’, H’ of all 8 words A, …, H that had been bound together. We needed to use here lower thresholds for the memory neurons so that 2 out of 8 words on the input layer can already activate sufficiently many of them.

**Fig. 3:**
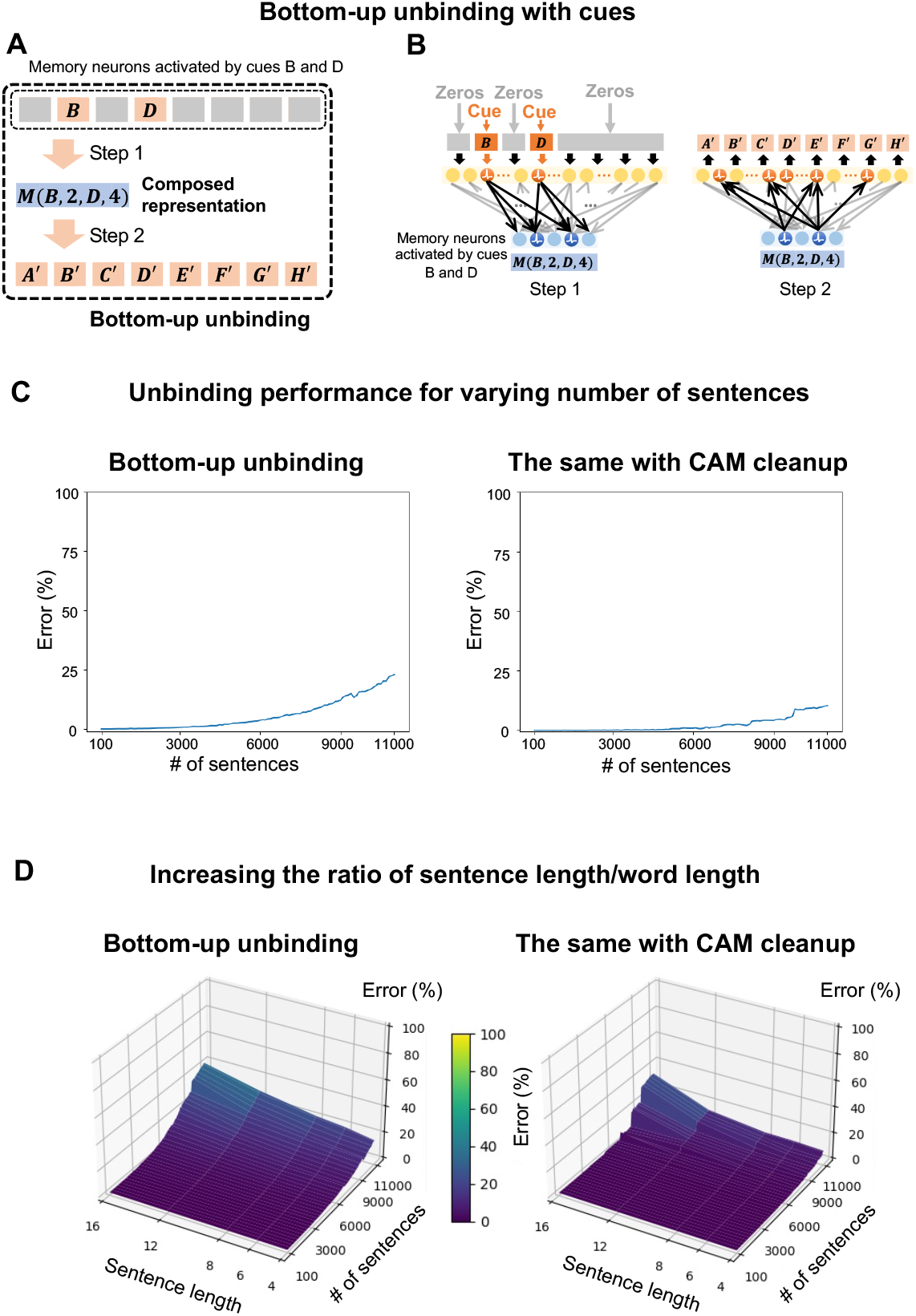
Bottom-up unbinding. **(A, B)** Scheme for bottom-up unbinding. Note that, unlike unbinding for currently existing HDC methods, the composed representation was not available for unbinding. Only 2 arbitrarily chosen words B and D, here in positions 2 and 4, were given as cues. **(C)** Performance of bottom-up binding with 2 out of 8 words as cues, with and without clean-up of decoded words via a BTSP-based CAM. The error measure is the same as in Fig, 2F. **(D)** We consider here the case where 2 randomly selected words from longer sentences with 12 or 16 words are used as cues. In other words, just 1/6 or 1/8 of the bits in the sentence are used as cue. Good performance can still be achieved in this case, but degrades gracefully when the number of sentences becomes very large.

The bottom-up unbinding performance is almost as good as in the previously considered case of top-down unbinding. Fig. 3D shows that an even smaller fraction of words than 1/4 suffices: Even when just 2 out of 12 or 16 words in a sentence are provided as cues, bottom-up unbinding works still well. Obviously, bottom-up unbinding is a direct consequence of the attractor feature of BTSP-based HDC. It has no analogue in previous HDC approaches.

We had considered here the case where for each pair of words only a single composed representation (sentence) had been presented that contained these two words at their respective positions. In other words, bottom-up unbinding had a unique solution. In the next section we consider the case where this is no longer guaranteed.

### 2.5 Performance comparison with Vector-Symbolic Architectures on a benchmark task

We consider here a benchmark task that has previously been considered in (Frady et al., 2020; Kent et al., 2020; Renner et al., 2024): Extracting information from high-D codes for a composed visual scene consisting of 3 colored letters at different positions in 2D, see Fig. 4A for an illustration and Methods for details. Whereas the VSA approach of (Renner et al., 2024) uses algebraic operations, component-wise addition and multiplication, for encoding a visual scene consisting of several letters, BTSP-based HDC generates a code by applying BTSP to an input vector whose “words” are encodings of individual letters, their color, and their position.

**Fig. 4:**
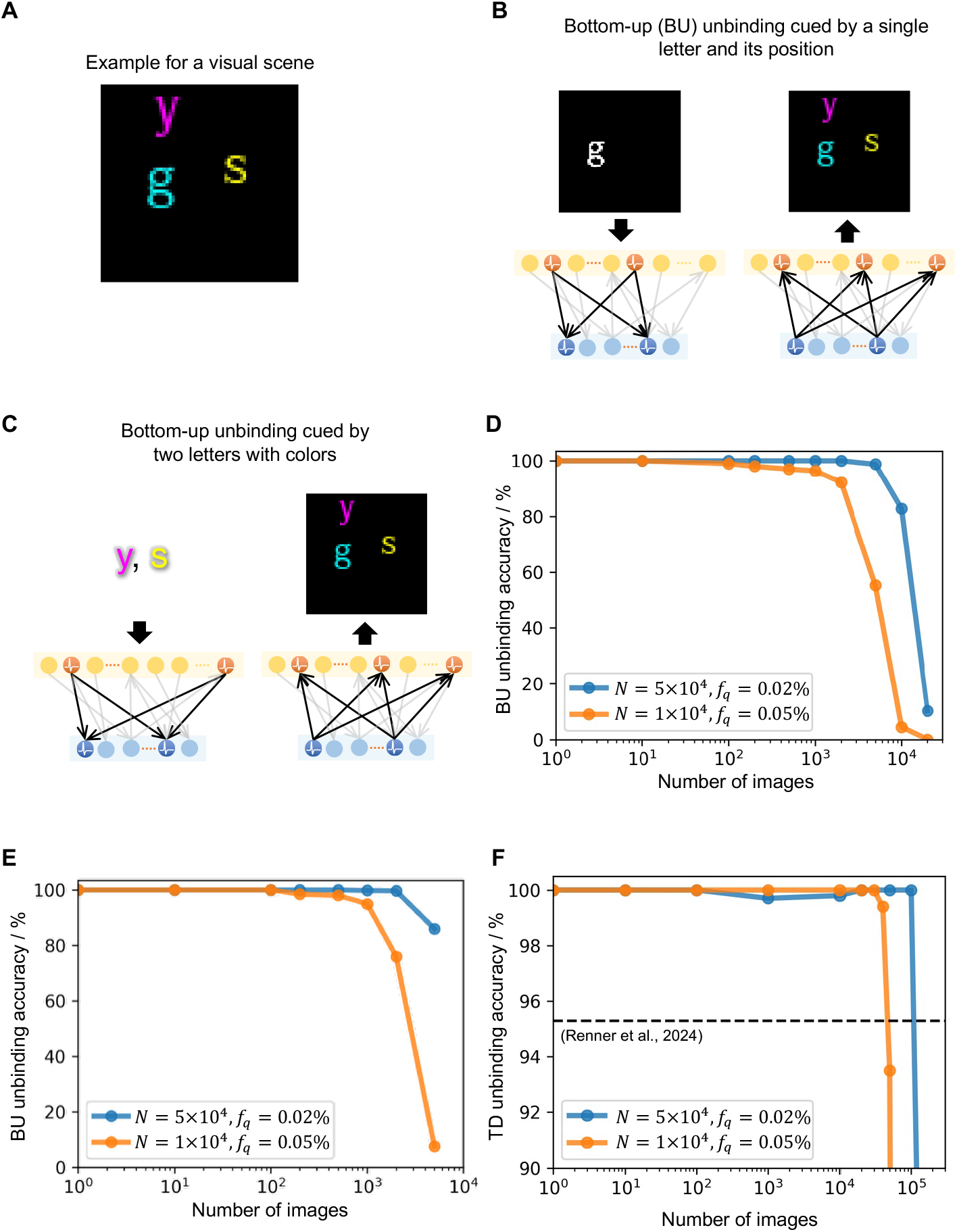
Comparison of BTSP-based and VSA-based unbinding. **(A)** An example visual scene contains three letters, each with one of seven possible colors and (*x, y*) coordinates within a 64 × 64 grid as features. **(B)** Information retrieval from BTSP-based binding that can not be carried out for VSA-binding: Retrieval of the full visual scene (shown on the right) by providing a single letter and its position as cue (shown on the left). The accuracy is shown in **(D). (C)** Retrieval of the full visual scene (shown on the right) by providing two letters and their color as cue (shown on the left). The accuracy is shown in **(E)**. Recover the full visual scene by providing two letters with their color. The accuracy is shown in **(E). (F)** Top-down (TD) unbinding accuracy for BTSP-based binding. This is shown as function of the total number of visual scenes that have been bound. The dashed horizontal line is the estimation of (Renner et al., 2024) for the accuracy of their approach (computed as 98.4%^3^ ≈ 95.3%). One sees that unbinding of BTSP-based binding works with much higher accuracy, unless the total number of bound representations is larger than the dimension *N* of the high-D vectors.

#### 2.5.1 Comparison on bottom-up unbinding

We first show in Fig. 4 B - E that a composed visual scene can be retrieved by just providing a single letter and its position as cue, or the identities of two letters without positional information. It is assumed here for simplicity that only a single scene fits to this cue, otherwise more complex retrieval operations are needed (see the next section). But also if the solution is unique, VSA-based binding cannot solve this task due to the non-invertability of the underlying algebraic operations. In particular, in VSA-based approaches also the composed representation has to be provided in addition to a component of the content. In contrast, Fig. 4 D, E show that bottom-up unbinding works very well for BTSP-based binding, without requiring presentation of the composed representation.

#### 2.5.2 Comparison on top-down unbinding

According to the reviews (Kleyko et al., 2022, 2023) unbinding is in classical HDC approaches only possible if besides the composed representation also a large fraction of the bound content is available. For example, in order to recover for a component of a high-D vector both factors x and y from their product, both the product and one of the two factors have to be given. However, it is shown in (Frady et al., 2020; Kent et al., 2020; Renner et al., 2024) that a specific type of recurrent neural network, a resonator network, can carry out a factorization of high-D vectors if a code-book is provided that lists all possible high-D vectors that could appear as factors. There is no theoretical guarantee for the convergence of a resonator network to a solution, but it usually does converge, as shown in these publications. For example, according to (Renner et al., 2024), a single letter is recovered for the benchmark task that we consider with probability 98.4%. In the following we compare the results for decoding of VSA-codes with a resonator network with results for decoding from BTSP-based binding with regard to accuracy, computational complexity, unbinding speed, requirements for a neuromorphic implementations, and scaling properties.

##### Accuracy

As mentioned above, the accuracy of recovering a single letter correctly from a composed VSA-representation is 98.4%. In order to recover all three letters for an instance of the benchmark task one has to subtract the code for the unbound letter from the composed representation, and then repeat the application of the resonator network. Hence all three letters can be correctly extracted with an accuracy equal to the 3rd power of 98.4%, i.e., with an accuracy of about 95%. In contrast, BTSP achieves for the same task an accuracy of 100%, see Fig. 4 F.

##### Computational complexity and speed of unbinding

Computations are carried out in both approaches on N-dimensional vectors, where N is typically relatively large (N = 10,000 in (Renner et al., 2024), N = 10,000 or 50,000 in our approach). The vectors in our approach are sparse binary vectors, where the density of 1’s is 0.05% or 0.02%, which is in the same range as the measured sparsity of neural codes in the brain (de Vries et al., 2020).

In contrast, the vectors on which computations are carried out in (Renner et al., 2024) are dense vectors with continuous-valued complex numbers as components (instead of bits, as in our approach). Besides iterated computations on these dense vectors, resonator networks require multiplication of matrices whose dimension scales with the size of the code-book, i.e., with the number of possible vectors that could potentially occur as high-D code for an item. In addition, whitening via singular value decomposition (SVD) is needed for the same high dimension. In contrast, BTSP unbinding does not involve matrix multiplication or SVD. It only requires multiplications of binary matrices (that hold binary synaptic weights) with sparse binary vectors. Furthermore, this unbinding can be carried out with a total of two parallel computation steps (see Fig. 9 A). These vector-matrix multiplications can be implemented very efficiently through in-memory computing by sending a vector of binary inputs into a memristor crossbar (Jung et al., 2022; Lanza et al., 2025). One further parallel computation step can be used for cleaning up the unbound items via a BTSP-based CAM (Fig. 9 B).

Top-down unbinding with resonator networks requires according to (Renner et al., 2024) to run the resonator network for about 30 steps in order to attain a high probability that it has converged to an item (letter) in the visual scene. This item is then subtracted from the composed representation, and the resonator repeats this computation on the remainder, until all items have been extracted. Hence unbinding for the benchmark task requires in total about 90 parallel computation steps of the resonator network, and in addition matrix multiplications and SVD for a dimension that is given by the size of the code-book. Furthermore, for each extracted item a Hopfield network (typically requiring around 20 steps until convergence) is applied in addition to clean up the code for the extracted item. Altogether this requires roughly 60 further parallel computing steps.

##### Requirements for a neuromorphic implementation

The main computation steps for BTSP-based binding and unbinding can be implemented through inmemory computing with memristor arrays, with parameters stored in simple memristors with at least two distinguishable conductance values. Note that all matrix and vector entries in this approach are just binary valued. The k-WTA operation for linear output neurons can also be carried out in most neuromorphic chips using lateral inhibition (Wang et al., 2019; Abdoli and Safari, 2020; Wang et al., 2023). If needed, one can actually eliminate the k-WTA operation by choosing a value for the firing threshold of output neurons so that the number of activated output neurons is in a desirable range. The most basic types of neuron models (Mc-Culloch Pitts neurons, linear neurons, single-compartment spiking neurons) suffice for that.

In contrast, the approach of (Renner et al., 2024) requires 4-compartment neurons. The entries of matrices that occur in this approach assume many different values, in fact a number of values that scales with the size of the code book for cleanup of decoded items via a Hopfield network. Hence these entries have to be stored in a less efficient way in a digital memory. In contrast, the in-memory computing approach for implementing BTSP-based binding and unbinding is likely to provide drastic advances in energy-consumption and latency.

##### Scaling properties

The unbinding approach of (Renner et al., 2024) works most efficiently if there are relatively few options for the identity of each item in the composed representation (there are just 26 for their benchmark task that we consider), since the number of possibilities for each item (the code book length) becomes a dimension of the matrices that need to be multiplied in their approach. This approach works most efficiently if the number of items in the scene remain small, since each additional item in a visual scene reduces the decoding accuracy by the factor 0.984. In contrast, BTSP-binding works also for very large numbers of possible items, and no code book has to be given in advance. Hence it can also be applied to visual scenes where the visual items are faces of people, and new faces can be added in an online manner. It can also be used for carrying out hierarchical binding, where composed representations on a lower level become tokens for higher level binding operations (see Section 2.7). Note that each item (component) in the preceding sections could be anything that can be encoded by a sparse binary vector of length 6000. We considered more specifically the case where there were on average around 30 1’s in these component vectors. The number of items which can be encoded in this way is obviously extremely large (for a precise estimate one needs to evaluate the combinatorial term 6000 choose 30, which yields a number larger than 10^80^). Also visual scenes with more than 3 items can be handled efficiently, without reducing the accuracy of unbinding in a significant manner (we had considered in the preceding sections sentences that consisted of 8 words). On the other hand, the total number of visual scenes can be large, but not arbitrarily large for the BTSP-based approach (it is in the VSA approach only implicitly constrained by the sizes of code books for items). For example, BTSP-based binding and unbinding works for the case of vectors with dimension N = 10,000 with 100% accuracy for M = 40,000 visual scenes, and for the case N = 50,000 for M = 100,000 visual scenes, see Fig. 4) F.

### 2.6 Bottom-up unbinding with ambiguous cues

We had assumed in the previously discussed bottom-up unbinding tests that just a single sentence had been learnt that was consistent with the given cue. We now drop this uniqueness assumption, and consider the case where there may exist several correct results for bottom-up unbinding. It is obviously not desirable that the network produces a superposition of them. Our brain tends to produce in this case different correct results of bottom-up unbinding in a sequential manner. We show that by adding an additional network module, a recurrently connected neural network, see Fig. 5A, our model can also do that. This recurrent neural network roughly corresponds to area CA3 in the brain, and for simplicity we refer to it as CA3 in our model. Corresponding to the conjecture role of area CA3 in the brain, we enable this recurrent neural network to act as associative memory that can store different correct input completions as attractors. It was shown in (Li et al., 2024) that recurrent synaptic connections within CA3 are quickly adapted by a Hebbian variant of BTSP. In this variant of the plasticity rule also postsynaptic firing is required, see Fig. 5B. Again, we consider the simplest case where these weights can only assume values 0 or 1. This plasticity rule is activated whenever one of the given set of input patterns (sentences) is encoded by the firing activity of the input neurons. More precisely, this synaptic plasticity within the recurrent network is applied after BTSP has been applied to the synapses from the input neurons. As a result, these activations of the recurrent network become attractors of the dynamics of the recurrent network.

**Fig. 5.**
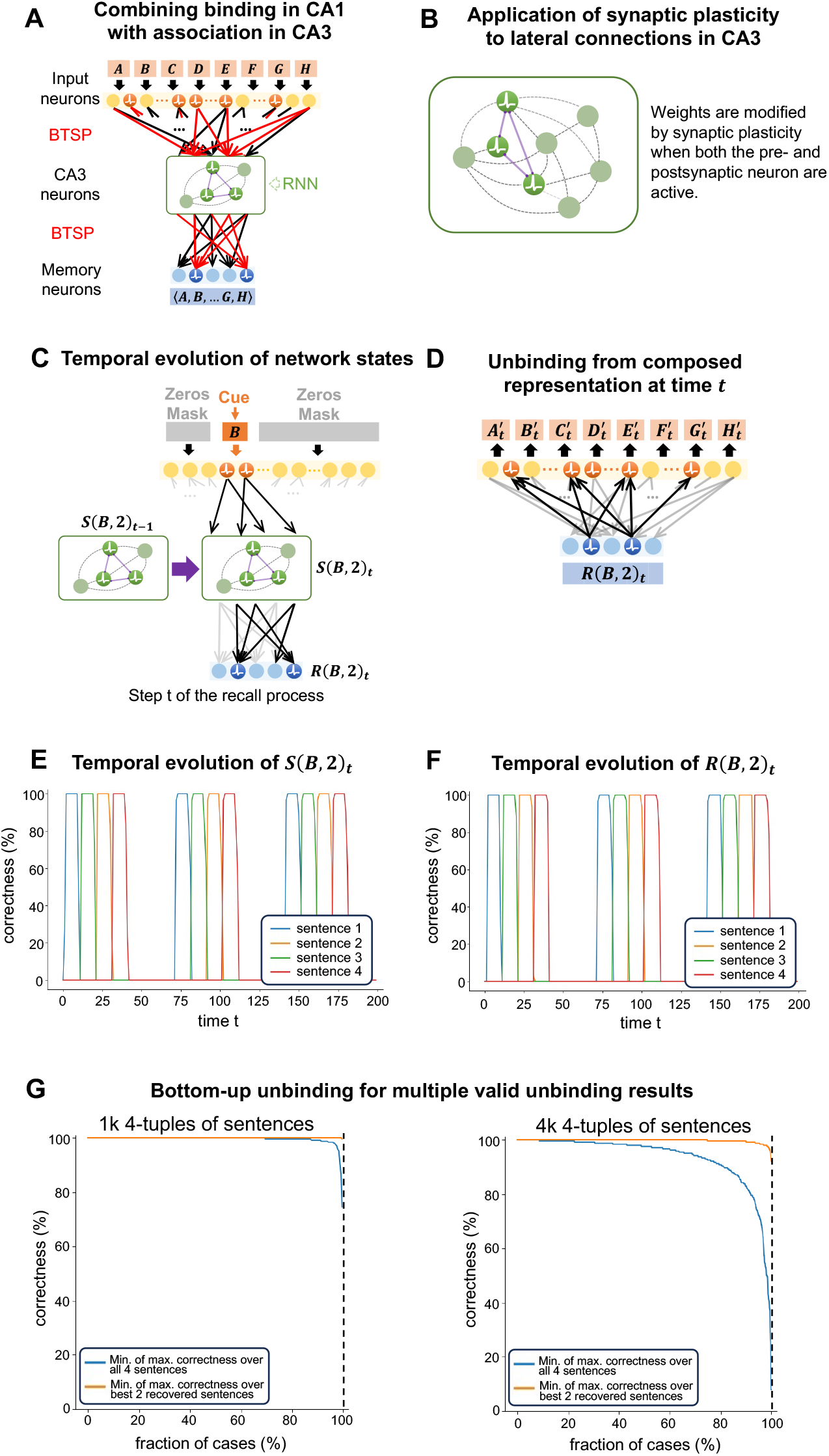
Bottom-up unbinding from single words when several correct unbinding results exist. **(A)** During the binding process, a newly added recurrent network module is positioned between the input and memory layers to enable a sequential dynamics of unbinding results. **(B)** Lateral connections within the recurrent network module are trained through a Hebbian variant of BTSP that was found in area CA3 of the brain (Li et al., 2024). **(C)** While a cue is presented on the input layer, the network state *S*(*B*, 2)_*t*_ of the recurrent network module evolves dynamically, where the next network state depends both on the cue and the preceding network state. These network states in the recurrent network module induce a sequences of activity patterns *R*(*B*, 2)_*t*_ of the memory neurons. **(D)** Each activity pattern *R*(*B*, 2)_*t*_ of the memory neurons induces via the feedback connections at each time step a possible result 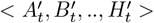 of bottom-up unbinding. **(E)** Evaluation of the similarity (measured via cosine) of the dynamically evolving network states *S*(*B*, 2)_*t*_ of the recurrent network module when the word B is presented as cue at the 2nd word position to its first state when a full sentence (one of the 4 possible solutions) is presented on the input layer. One sees that the network state cycles repeatedly through the 4 states (attractors) that result when one of the 4 possible solutions of this bottom-up unbinding test is presented on the input layer. **(F)** The same for activity patterns of the memory neurons. One also sees there repeated cycling through activity patterns that result as first activity pattern when one of the 4 possible unbinding solutions (i.e., full sentences) are presented on the input layer. Note that we have in our model no lateral connections between neurons on the memory layer (reflecting low lateral connectivity of pyramidal cells in area CA1). Hence the memory neurons would not be able to produce a sequence of different possible solutions with the preceding recurrrent network module. **(G)** Evaluation of the sequence of unbinding results that are produced in the input layer through feedback connections as indicated in panel D, both for the case of a total number of 1000 sentences, and for the case of 4000 sentences. The blue curve denotes the fraction of ambiguous single word cues for which each of the 4 possible correct unbinding results (sentences) were reproduced at some time point with the error indicated on the y-axis. The yellow curve shows for which fraction of them at least 2 of the 4 possible solutions were reproduced with the error indicated on the y-axis.

One can avoid that the dynamics of this recurrent network module remains in a single attractor by adding an adaptation property to its neurons. This adaptation property makes each neuron reluctant to fire continuously for a large number of time steps (see (Chen et al., 2022) for models for adapting biological neurons and (Rao et al., 2022) for applications in neuromorphic hardware). With these neuron models the recurrent network moves sequentially through all attractors that are consistent with the current network input. In the illustration in Fig. 5C the activity of this recurrent module at time t for the cue word B at the 2nd word position is denoted by *S*(*B*, 2)_*t*_. Synaptic connections to memory neurons transforms this code into another neural code *R*(*B*, 2)_*t*_. When the recurrent network module moves through its attractors, the memory layer moves through corresponding codes *R*(*B*, 2)_*t*_. Each of them can be decoded by the feedback connections into a possible sentence 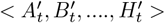 as indicated in Fig. 5D.

We demonstrate the functionality of this method by considering the case where one has for each word in a particular position 4 learned sentences that are consistent with this cue. When a single word is given as cue, for example in Fig. 5C the word B in the 2nd word position, neural activity in the recurrent network module cycles through 4 attractors (Fig. 5E), where each of these attractors had been induced during learning of lateral weights by one of the 4 possible sentence completions of this cue. This dynamics in CA3 induces a corresponding cycling through 4 composed representations *R*(*B*, 2)_*t*_ in the memory neurons (Fig. 5F). The correctness values on the y-axis in Fig. 5 E and F (see Methods for details) result by comparing the neural activity over time during the presentation of the single word as cue with the activity induced by presentations of the 4 possible sentence completions as network input. One sees, that the network activity for the single word as cue cycles between those that are induced by the 4 sentences.

Each activity pattern on the memory layer induces in the input layer through the feedback connections (Fig. 5D) a proposition for bottom-up unbinding. The quality of this unbinding result is shown in Fig. 5G for two cases where 1000 and 4000 sentences had been learnt. For each of the 4-tuples of sentences that we used in our experiments as correct completions of an input cue we measured the maximal correctness of unbinding over 200 time steps, for each of the 4 possible sentence completions (taking the minimum over them). Hence a value y for the min. of max. correctness over all 4 sentences in a 4-tuple of correct unbinding results indicates that each of the 4 possible sentence completions was reproduced at some time point on the input layer at least with correctness y. The x-axis in Fig. 5G indicates for which fraction of 4-tuples of sentences this value y could be achieved. For example, in the case where one has 1000 4-tuples of sentences, for about 90% of them a value of y close to 100% (i.e., perfect decoding for each of them) could be achieved.

The yellow curves in Fig. 5G give corresponding measurements if one is satisfied to recover at least 2 of the 4 possible sentence completions at some time point.

Although unbinding from ambiguous cues is a natural real-world challenge, which is mastered quite well by our brain, there exist to the best of our knowledge no results for unbinding from ambiguous cues for standard HDC binding.

### 2.7 Iterated binding and unbinding

A hallmark of any neural network model for compositional computing is the ability to iterate compositional processes, e.g., to form sentences from words and then stories from sentences, as indicated in Fig. 6A. While this scheme only addresses the case of hierarchical binding, it is also desirable to expand a composed representation by adding further content or structural information in an online manner, as indicated in Fig. 7A. The real challenge for iterated binding is of course the capability to be able to fully reverse it through iterated unbinding (see Fig. 6B and Fig. 7B). One concern on the technical level is a likely accumulation of errors at different stages of iterated unbinding. But we show in Fig. 6 C,D that the performance of iterated top-down unbinding is quite good for hierarchical binding through BTSP. Furthermore, it can be significantly improved by using CAM cleanup. The results for iterated online binding are similar, see Fig. 7C. For simplicity we refer to the items that are bound together as words, and to the lists of words that are bound together through iterated binding as a sentence. But in an application to natural language the latter could also represent a story that consists of several sentences.

**Fig. 6:**
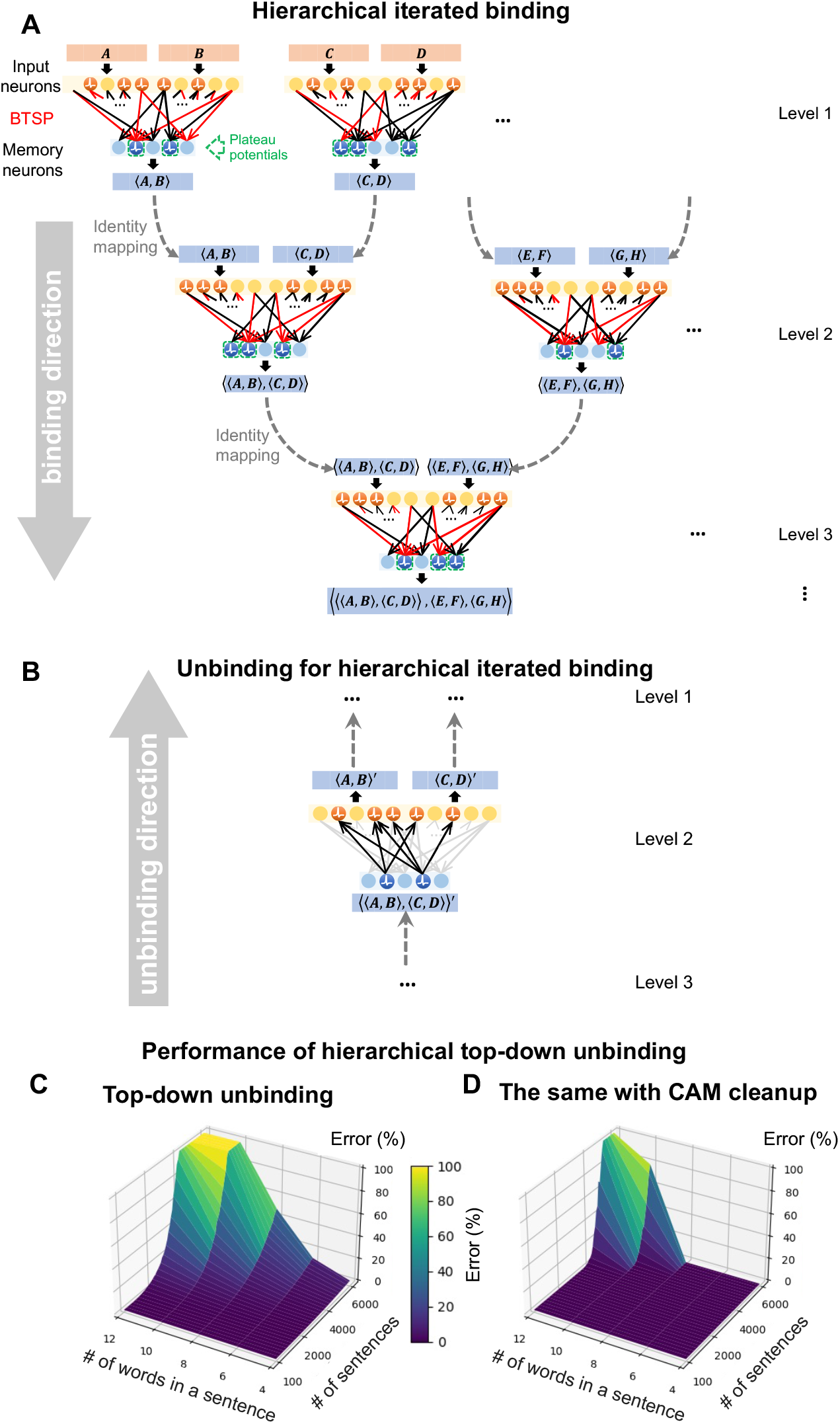
Hierarchical iterated binding and unbinding. **(A)** Scheme for hierarchical iterated binding. **(B)** Scheme for hierarchical top-down unbinding. **(C**,**D)** Performance of top-down unbinding for iterated hierarchical binding, with and without a BTSP-based CAM for cleaning up the words that are top-down generated at the input layer. The CAM significantly improves the unbinding result.

**Fig. 7:**
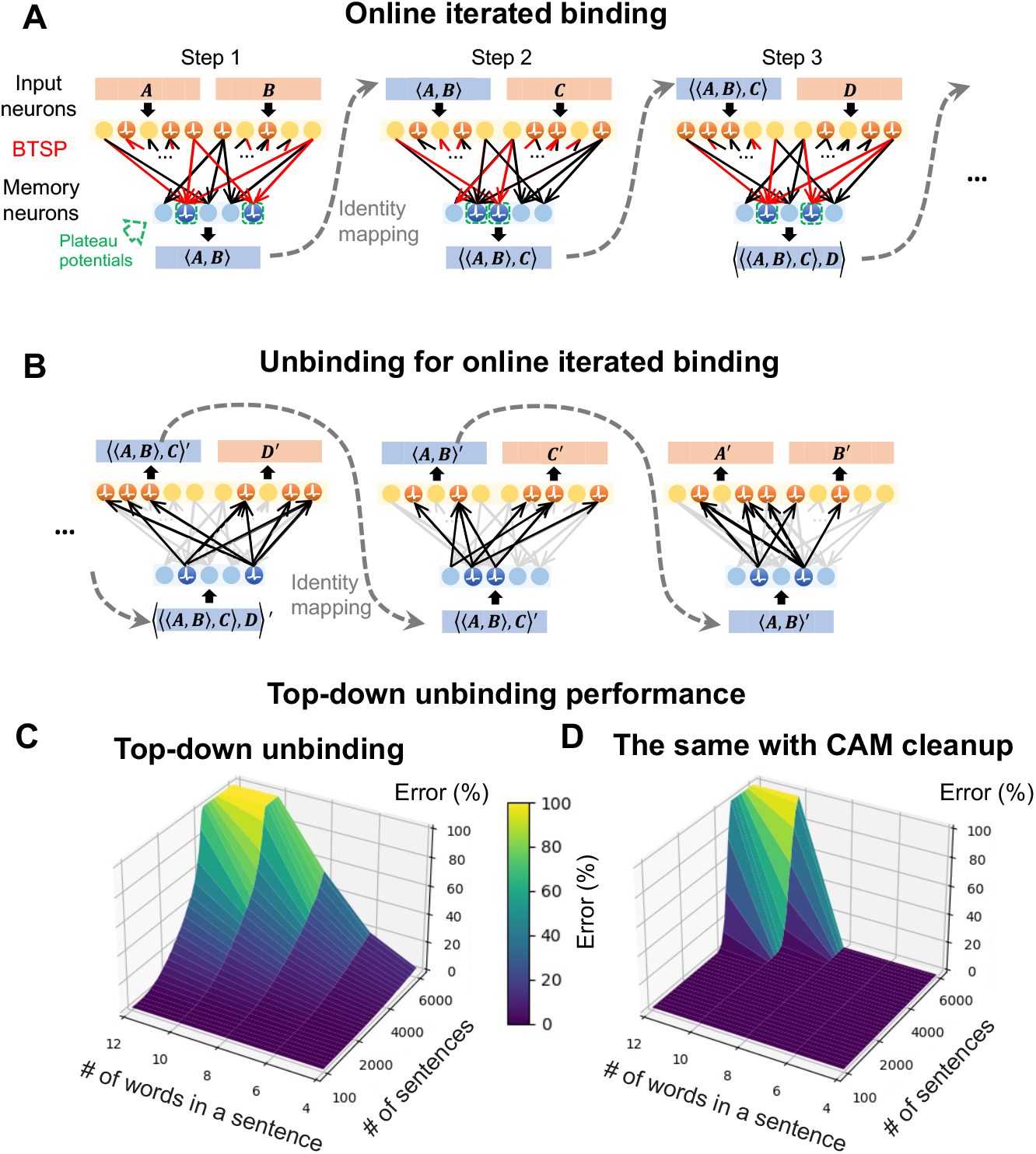
Online (linear) iterated binding and unbinding. **(A)** Scheme for online iterated binding. **(B)** Scheme for online iterated unbinding. **(C**,**D)** Performance of top-down unbinding for iterated online binding, with and without a BTSP-based CAM for cleaning up the words that are top-down generated at the input layer.

Note that there are no comparison results for iterated top-down unbinding for classical HDC, since top-down unbinding is not possible there.

### 2.8 Advancing the assembly calculus for modeling language processing in the brain through BTSP-binding

Binding operations play a central role for models of higher-level brain computations, especially for language processing, based on an assembly calculus (Papadimitriou et al., 2020; Müller et al., 2020). The idea of these models is that assemblies of neurons, such as for example an assembly of concept cells for a particular concept or structural information, are tokens for brain computations.

Complex computations and data structures for semantic content and structural annotations can be formed with these tokens if there is a mechanism for binding two content tokens together to form another assembly. Obviously, a binding mechanism is needed that is able to bind assemblies together very fast, when a particular episode is experienced or a particular sentence is heard. Hence synaptic plasticity mechanisms such as STDP, where dozens of iterations of a learning trial are needed for changing a synaptic weight (Froemke et al., 2010), are less plausible as biological implementation of binding. For natural language processing, and also for the frequently postulated internal language of thought of the brain (Kazanina and Poeppel, 2023), one needs in addition the capability to bind an assembly that represents a concept, such as “cat”, to an assembly that indicates a specific role for that concept in a sentence or episode, such as “cat as agent” (Frankland and Greene, 2020b), see Fig. 8A. Previous models (Papadimitriou et al., 2020; Müller et al., 2020) proposed that this is carried out via an instantaneous “projection” of an assembly into another cortical area that represents specific semantic roles. Obviously, binding through BTSP offers itself as a suitable synaptic plasticity mechanism for that.

**Fig. 8:**
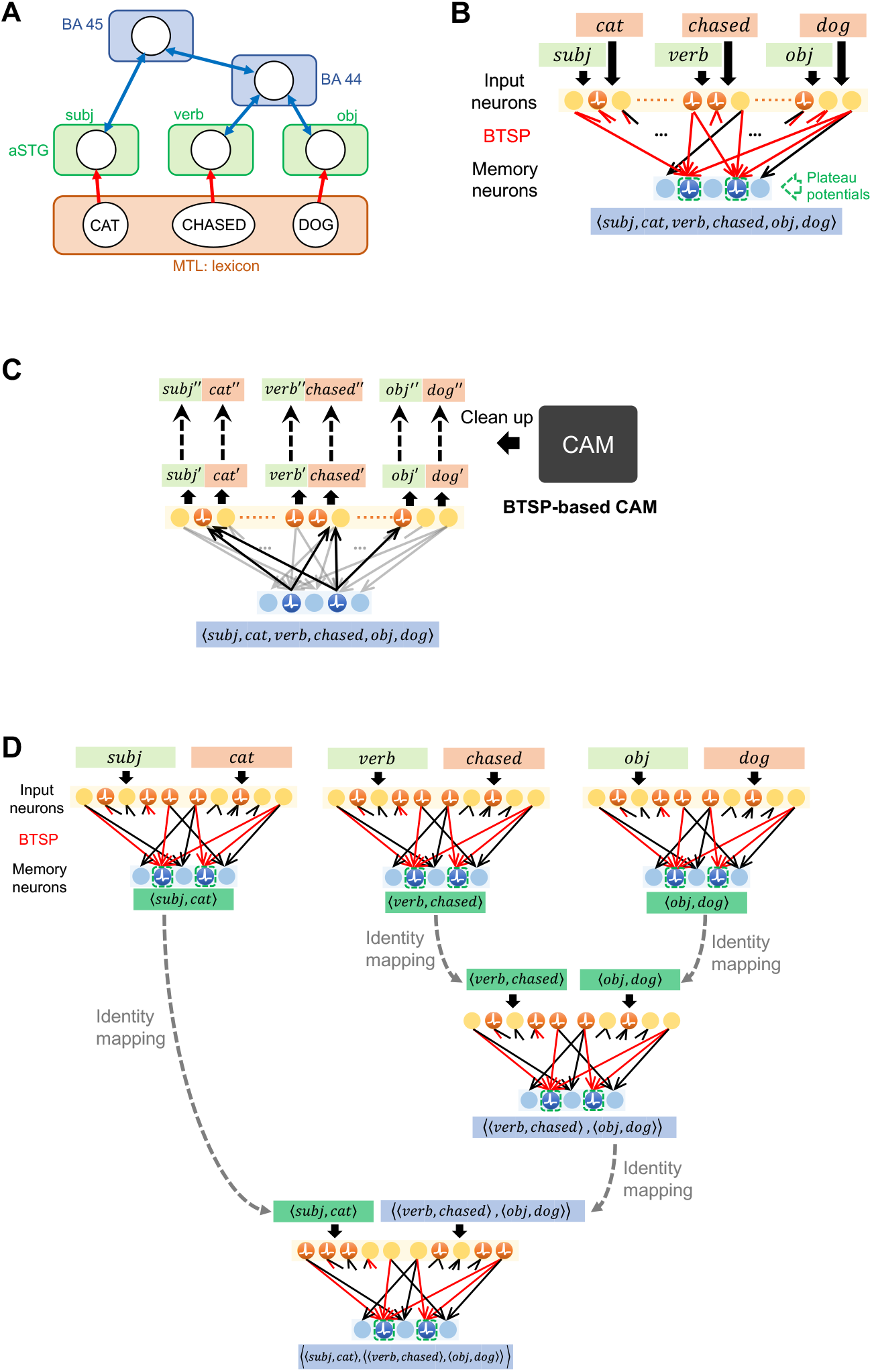
Application of BTSP-based binding in models for language processing in the brain. **(A)** Scheme for syntactic analysis of a sentence through interaction of several cortical areas (MTL, sSTG, BA44, BA45) proposed by (Papadimitriou and Friederici, 2022) (reprinted from this publication). **(B)** Binding of the 3 words from panel A and of their syntactic roles into a composed representation through a single application of BTSP, see also Fig. 2A. **(C)** Decoding of constituent words and their syntactic roles from an internal composed representation of the sentence, as generated through BTSP in panel B. As in Fig. 2E, a BTSP-based CAM can be used to clean up the decoded components. **(D)** Hierarchical model for producing a composed representation of the same sentence through BTSP-binding as in panel A, as special case of Fig. 6.

**Fig. 9:**
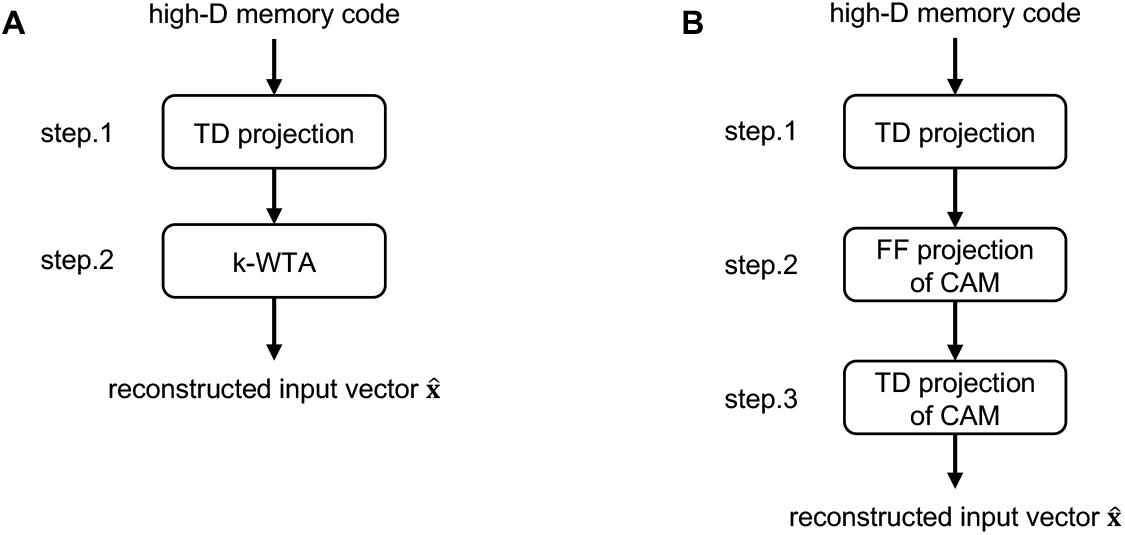
Computation steps of TD unbinding for BTSP-based binding. **(A)** In the most basic form, as in Fig. 4, see Sec. 4.4), the model requires only two computation steps: TD projection and a k-winner takes all (WTA). **(B)** If one uses in addition a CAM for cleanup of codes for retrieved components (as in our other figures), the model requires three computation steps. TD projection, feedforward (FF) projection of CAM, and TD projection of CAM.

**Fig. 10:**
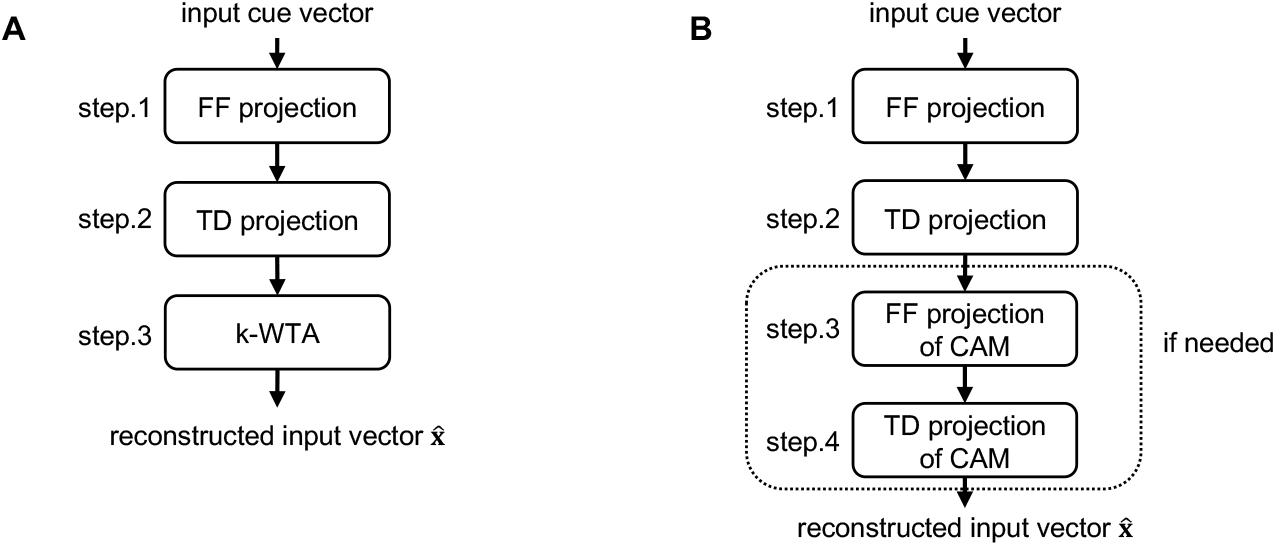
Computing steps of BU unbinding (only possible for BTSP-based binding). **(A)** We first consider the case without CAM (in Fig. 4, see Sec. 4.4). Here the BU unbinding requires three steps. **(B)** With a CAM used in BU unbinding (as in our other figures). one needs up to four steps.

Binding of two assemblies by strengthening synaptic connections between neurons in the two assemblies through STDP has been considered in previous work (Papadimitriou et al., 2020; Müller et al., 2020). But, the experimental data on instantaneous formation of conjunctive memories in area CA1 (Bittner et al., 2015; Zhao et al., 2022) suggest a different solution. There not the synapses between concept cells for different concepts are modified, but direct or indirect synaptic connections between concept cells and neurons in another area, as indicated in the scheme of Fig. 8B. The general compositional computing operations that are supported by BTSP according to Fig. 2 and 3 can readily be used as basic operations of such an alternative brain calculus, see Fig. B and C. Feedback connections and larger loops, such as the big-loop recurrence between CA1 and cortical areas (Koster et al., 2018), allow to iterate and reverse such binding process, see Fig. 8D for an illustration. Binding through BTSP can handle such iterated binding and unbinding quite well, as shown in Fig. 6 and 6. Hence, we propose to improve the brain calculus of (Papadimitriou et al., 2020) by employing binding through BTSP via composed representations that act as pointers to the assembly codes. This model is supported by the recent experimental data of (Kolibius et al., 2023).

## 3 Discussion

Compositional computing, where a few items from a limited repertoire of items are bound into a myriad of different composed representations that acquire new meaning not present in the items, is a key factor of computational intelligence and intelligent communication. But an implementation of binding with flexible retrieval possibilities is a fundamental problem if items are represented in a distributed manner, as is the case for brains (Frankland and Greene, 2020a; Kurth-Nelson et al., 2023; Kazanina and Poeppel, 2023; Piantadosi et al., 2024) and most current AI approaches.

The binding problem has been centrally addressed in work on HDC (Kleyko et al., 2022, 2023). Previous HDC approaches, such as VSAs and semantic pointers, employed algebraic operations on high-D vectors for binding. We have shown here that information retrieval capabilities can be substantially enhanced if one uses instead a mechanism that the brain employs for binding, BTSP (Bittner et al., 2015; Zhao et al., 2022). A simple model for BTSP from (Wu and Maass, 2025) suffices for that. It enables HDC to carry out top-down unbinding, i.e., retrieval of all tokens that have been combined in a composed representation (see Fig. 2), with very small delay. This top-down unbinding can be carried out very efficiently even in the case of iterated binding, where composed representations become tokens for binding on a higher level (Fig. 6, 7, 8). Furthermore, the attractor features of composed representations that BTSP provides also make these operations robust against errors that may arise in implementations on highly energy-efficient analog hardware such as memristor crossbars (Fig. 2F). In contrast, top-down unbinding is in general impossible for binding approaches that are based on algebraic operations, such as VSAs and semantic pointers. However, resonator networks can do that, provided they get a codebook for all possible tokens that might occur (Frady et al., 2020; Kent et al., 2020; Renner et al., 2024). We have shown that in comparison with resonator networks applied to VSA-binding, top-down unbinding is for BTSP-based binding substantially faster, simpler to implement, and more accurate (see Fig. 4F). Furthermore, it can also be applied if the number of possible tokens is very large, as for example in the case of iterated binding, where composed representations on a lower level become tokens for binding on a higher level. BTSP-based binding also offers new information retrieval options that were not supported by any previously proposed binding mechanism. An example is bottom-up unbinding, where only a minor fraction of bound tokens are provided as cues, and not the composed representation (Fig. 3). We have also shown that if the cues are ambiguous, different solutions can be retrieved sequentially when an CA3-like attractor module is added (Fig. 5).

BTSP-based binding provides a new method for modeling the neural implementation of a language of thought (Kazanina and Poeppel, 2023) and of natural language processing in the brain (Fig. 8). In particular, the Merge operation, which has been proposed in linguistic theory as universal grammatical binding operation (Chomsky et al., 2023), can easily be modeled with BTSP-based binding, thereby enhancing the models of (Papadimitriou and Friederici, 2022; Liu et al., 2023). Furthermore, BTSP provides a method how assemblies of neurons can instantly be bound together, rather than requiring numerous iterations as in previous models (Müller et al., 2020; Papadimitriou et al., 2020).

Binding through BTSP only requires binary weights. This makes our models consistent with the rather limited number of weight values that biological synapses can assume (Bartol Jr et al., 2015), and with constraints of especially efficient hardware implementations of synaptic weights in memristors that can only assume a small number of conductances (Wang et al., 2020; Khaddam-Aljameh et al., 2022). In addition, in contrast to most neural network models, BTSP-enhanced HDC operates with sparse neural codes, both for content items and composed representations. Therefore it can be implemented in the most energy-efficient activity regime, both on spiking and non-spiking neuromorphic hardware (Davies et al., 2021). A promising direction for future research is a combination of BTSP-enhanced HDC with trained operations on high-D vectors with in DNNs (deep neural networks) or LLMs, for example in order to enhance analogical reasoning capabilities of energy-efficient neuromorphic hardware (Hersche et al., 2023; Camposampiero et al., 2025).

## 4 Methods

**Table 1:**
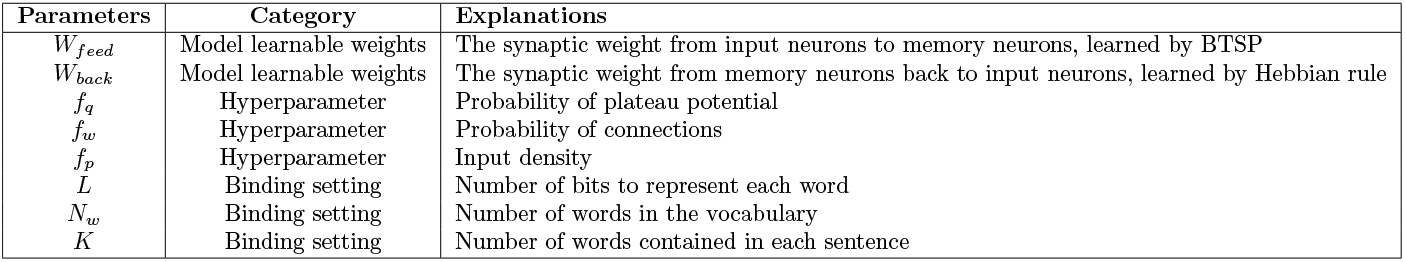
Definitions and explanations of key parameters.

### 4.1 Implementation of BTSP

#### 4.1.1 Details to the BTSP rule

We use throughout the simpler version of the BTSP rule (called core BTSP in (Wu and Maass, 2025)), in which the explicit probabilistic factor (set as 0.5) governing synaptic weight updates is removed. It was shown in Fig. S1 of (Wu and Maass, 2025) that this simpler version produces essentially the same learning results as the rule with probabilistic factors that matches more closely the biological data (this factor arises from the uncertainty whether a synaptic input falls into the LTP or LTD part of the plasticity window, and the fact that LTP can only be applied to a binary weight if it has value 0, and LTD only if it has value 1).

Note however, that also this simplified BTSP rule maintains a stochastic element—since the gating signals (plateau potentials) are assumed to occur randomly.

In practical simulations, this simplified version enables replacing the original iterative sampleby-sample computation with batch-wise matrix multiplication (detailed in Sec. 4.1.2), thereby significantly enhancing analytical tractability and computational efficiency.

#### 4.1.2 Details to batch-wise BTSP learning

We show here how the BTSP rule can be implemented in parallel computations, enabling batch-wise simulations for accelerated processing. Let **x** = (**x**_1_, …, **x**_*n*_) denote binary input vectors representing presynaptic activity for different items, and let **q** = (**q**_1_, …, **q**_*n*_) denote synaptic plasticity windows corresponding one-to-one with these inputs, where the memory neurons selected within the synaptic plasticity windows will follow the update formula of Eq. (1). We consider a single update at iteration *k* for the synaptic weights **w**^(*k*)^ using the pair (**x**_*k*_, **q**_*k*_) under the BTSP rule described in Eq. (1). Specifically, focusing on the *i*^*th*^ input neuron (**x**_*k*_)_*i*_, the plasticity of *j*^*th*^ memory neuron (**q**_*k*_)_*j*_, and a single synapse (**w**^(*k*)^)_*ij*_—representing the connection from the *i*^*th*^ presynaptic neuron to the *j*^*th*^ postsynaptic neuron—the weight update rule can be expressed as:

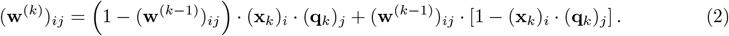

This equation can be generalized and compactly represented across all synaptic connections as:

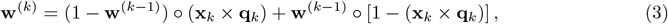

where ◦ denotes the element-wise product. For further simplification, we introduce modular arithmetic to equivalently represent the binary flipping operation (0 ↔ 1):

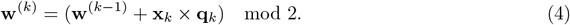

Thus, after *n* iterations, the weight matrix **w** can be concisely determined as:

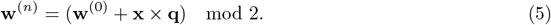

### 4.2 Details of binding and top-down unbinding (Fig. 2)

Given a sentence represented by binary atomic vectors *A, B*, …, *G, H*, each of length *L*, the binding phase employs one-shot learning via BTSP to train the feedforward synaptic weight matrix *W*_feed_, and to produce with these modified weights a composed representation ⟨*A, B*, …, *G, H* ⟩. Concurrently, direct feedback synaptic connections from memory neurons to input neurons are trained using a oneshot Hebbian-like learning rule to facilitate the unbinding process—recovering the atomic vectors *A*^*′*^, *B*^*′*^, …, *G*^*′*^, *H*^*′*^ from the composed representation. Specifically, if both presynaptic and postsynaptic neurons of a feedback connection are simultaneously active (i.e., both assume a binary value of 1), the synaptic weight of this feedback connection is incremented by 1; otherwise, the synaptic weight remains unchanged. If the weight value exceeded 1, it was set to a saturation value of 1. The feedback synaptic connections are adjusted only once for each network input, concurrently with BTSP. Parameter adjustments in the model are performed exclusively during the binding phase. Unless otherwise noted, in all of our experiments, we construct the vocabulary using 1000 words (*N*_*w*_ = 1000), initialize both the feedforward (*W*_feed_) and backward (*W*_back_) weight matrices with 0’s, and use default parameters for modeling: *f*_*p*_ = 0.005, *f*_*q*_ = 0.0025, and *f*_*w*_ = 0.6, consistently applied across both the binding model and the BTSP-based CAM. The CAM model is used to perform cleanup operations on restored words, which involves inputting the restored word into the CAM to activate the corresponding code in the CAM’s memory layer, followed by passing through the CAM’s feedback to restore it to the expected word. The clean-up process serves to filter out noise on restored words.

In Fig. 2F, top-down unbinding results based on sentences containing 8 words each (*K* = 8) are depicted for a range of sentence counts from 100 to 11,000, with intervals of 100 for each data point. Each word is defined by a vector of 6000 bits (*L* = 6000), consistently containing 30 active bits (*f*_*p*_ = 0.005). The size of the input layer is set to *K* times the word length, totaling 48,000, while the memory layer is sized at 12,000. During the binding phase, the threshold for memory neurons is set at 96, meaning neurons with activation levels of 96 or higher will spike, while those below will not. During the unbinding phase, the threshold for the input layer is determined using a grid search for each data point. For the BTSP-based CAM, the size of the input layer is equal to the word size, 6000, and the memory size is set at 12,000. The threshold for memory neurons is set at 12, and the threshold for the input layer for output is similarly obtained through grid search. It is important to note that the grid search process for threshold determination is conducted separately from the binding performance tests; optimal threshold values are determined using a different vocabulary prior to the binding experiments, and these predetermined thresholds are then fixed for use in the experiments. The primary metric employed is the percentage of incorrect bits, calculated as the ratio of the number of error bits to expected number of 1’s.

### 4.3 Details of bottom-up unbinding (Fig. 3)

The bottom-up unbinding evaluates the capability to recover original inputs when provided with only a subset of cues, while the remaining inputs are masked as zeros. Unless specifically stated otherwise, during testing of bottom-up unbinding, we default to randomly selecting 2 words from the sentence as cues. There are no restrictions on the positions of these cues; they can be chosen randomly. However, we ensure that the selected cues do not lead to multiple viable sentences. Cases involving ambiguous cues are discussed separately in Sec. 4.5. During the bottom-up unbinding process, we reduce the threshold in the memory layer to accommodate the decreased input intensity caused by the mask, ensuring the memory neurons can activate as expected. The reduction of the threshold involves a grid search to select the optimal solution suitable for the length and the number of sentences.

In Fig. 3C, bottom-up unbinding results based on 2 cues from 8-word sentences are depicted for a range of sentence counts from 100 to 11,000, with intervals of 100 for each data point. The experimental setup for binding is consistent with that described in Fig. 2F; we employ the same model used in top-down unbinding testing for each data point.

In Fig. 3D, bottom-up unbinding results are depicted for varying sentence lengths and counts. In this experiment, the sizes of both the input and memory layers are fixed at 48,000 and 12,000, respectively. The word length (*L*) is scaled according to the number of binding words (*K*), such that *L* = 48000*/K*. Correspondingly, the input dimension of the CAM is adjusted to the corresponding *L*, while the memory size is fixed at 24,000. A grid search for threshold reduction is applied for each data point, where we reselect the most suitable new threshold for the bottom-up process.

### 4.4 Details of our performance comparison with the HDC approach (Fig. 4)

We consider the same visual scene analysis task as considered in (Frady et al., 2020; Kent et al., 2020; Renner et al., 2024). Each visual scene consists here of 3 letters, each having one of 7 colors, and an arbitrary location of its center in a 64 x 64 pixel grid. In our implementation with BTSP-based binding each letter is encoded by the following bit string:

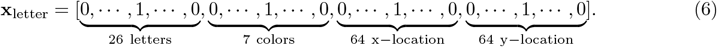

Each image is the concatenation of three letters (bit strings) of this type:

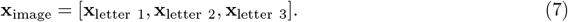

A visual scene was encoded in (Renner et al., 2024) somewhat differently. They used a sum of labeled pixels, rather than a standard input format for binding of a small set of tokens and their features, as for example the previously described input presentation. Hence their decoding task included an image segmentation task. But this image segmentation is trivial in the case which they considered, where images consist of 3 non-overlapping letter templates of fixed size and orientation. Hence a transformation of the input considered in (Renner et al., 2024) to the input format that we are using is trivial from a computational perspective. It can for example be carried out by a shallow ANN. Actually, such a transformation is also readily available for substantially more complex visual scenes, see e.g. (Spoerer et al., 2017).

*N* CA1 neurons were used in our implementation. Each neuron had a probability *f*_*q*_ to receive a plateau potential. All learning rules are also kept in its basic form as in Eq. (1) for the forward connection, and Hebbian with weight saturation at 1 for the backward connection. Specifically, if both presynaptic and postsynaptic neurons of a feedback connection are simultaneously active (i.e., both assume a binary value of 1), the synaptic weight of this feedback connection was incremented by 1; otherwise, the synaptic weight remained unchanged. If the weight value exceeded 1, it was set to a saturation value of 1.

#### Top-down unbinding

During TD unbinding, a learned image representation is fed into the network, and we evaluated if the network can reconstruct the same representation again.

Forward path is a simple fully-connected layer (with *f*_*w*_ = 1):

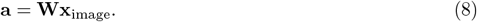

A threshold is applied onto the memory neurons, making the outputs binary:

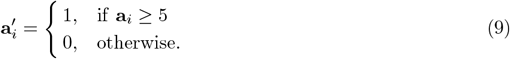

We choose the threshold value to be 5, which is an empirical choice. This task contains 12 non-zero inputs for each image (3 letters × 4 ones each). However, some synapses might be “forgot” after went through LTD. Therefore the threshold should be smaller than 12. Too large a threshold reduces the number of neurons that are activated by the input image, while too small a threshold may not able to shut down interfering neurons. The performance is relatively stable for a threshold in the range from 3 to 8.

Then, the backward path is also a linear layer:

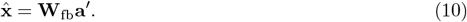

As before, we applied local-winner-takes-all:

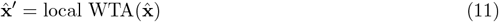

to clear up the reconstructed output.

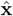 is the concatenation of three letters 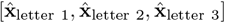 . For each of them, we applied:

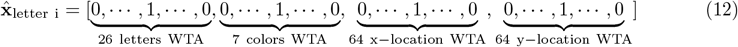

to decode a valid data information.

Then we compare if the reconstructed result 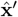 equals to the raw input image’s information vector **x**_image_, and report the reconstruction accuracy. Only a full match is considered as a success. We statistically count how many input images’ corresponding code can be decoded accurately.

#### Bottom-up unbinding

During data generation, we guaranteed that for each BU unbinding task, only one correct answer exist. For the first task cued by a single letter with its position information. The unique answer constrains the dataset size to (#letters×#x location × #y location)*/*3 = 35, 498. For the second task cued by two letters with their color information. The unique answer constrain limits the dataset size to 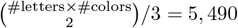.

The cue input has value 1 on the known information and value 0 on other information. For example, if one cues with a letter with its position (no color), the input representation of this letter is:

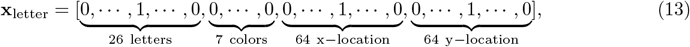

where the color input contains all zeros. We call now the input **x**_cue_, which corresponds to an identical image input **x**_image_.

Besides for the known parts that these two vectors are equal on corresponding dimensions. All other dimensions in **x**_cue_ are zeros.

Same as before, this input went through the forward layer (threshold by 3 for the single letter with position cue, and threshold by 4 for the two letters with colors cue). After thresholding, the memory neuron’s binary activation then went through the backward connections. Eventually, applying the same local WTA as in Eq. (12), we got a reconstructed output 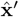.

We compare if 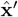 equals to the corresponding complete input vector **x**_image_. Only a full match is considered as a success. We statistically count how many cues can reconstruct the corresponding complete input accurately.

### 4.5 Details of bottom-up unbinding with ambiguous cues (Fig. 5)

We note that during recall via bottom-up unbinding, there can be cases where a set of cues corresponds to multiple valid sentences in memory. We refer to such cues as ambiguous cues, which evoke a superposition of all viable sentences stored in memory. To address this ambiguity, we introduce CA3 as an intermediate layer to temporally decouple the superposition, enabling the network to oscillate sequentially among all valid patterns. Specifically, dense lateral connections and global k-Winners-Take-All (k-WTA) operations are incorporated within CA3, allowing attractors to form before propagating activation to the memory layer. We employ k-WTA to control the number of spikes in CA3, ensuring stability in the recurrent process, selecting the neurons with the highest activation levels to fire spikes while the rest remain inactive. Similar performance can be achieved without WTA competition if the neuron thresholds are properly adjusted to the level of activation that changes with iterations in CA3. One can use a grid search to find the appropriate thresholds for CA3 to achieve the desired number of one bits. The learning of weights for lateral connections within CA3 follows a Hebbian-type one-shot plasticity rule, similar as the one discussed in (Li et al., 2024). Synaptic weights, represented as *w*_*ij*_ connecting neuron *i* to neuron *j*, are binary and updated as follows: if both neuron *i* and *j* are simultaneously active, the synaptic weight is toggled between 0 to 1:

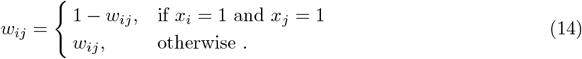

These weights were initialized as 0.

The schedule of steps where neuron outputs and synaptic weights are updated is as follows (see Algorithm 1 in the Supplement for further details):

Step 1. A sentence is presented as network input.

Step 2. BTSP is applied to the connections from input neurons to neurons in CA3.

Step 3. Neurons in CA3 are activated.

Step 4. BTSP is applied to weights from neurons in CA3 to neurons in the memory layer.

Step 5. Neurons in the memory layer are activated.

Step 6. Weights of recurrent connections in CA3 are subject to synaptic plasticity with the activation pattern from Step 3.

During testing, we conducted bottom-up unbinding experiments by presenting a single cue (among a total of 8 words per sentence) for the case where this cue appears at the same position in 4 different sentences for which the network was trained. Recurrent synaptic connections are activated in CA3, so that the network input generates a dynamic evolution of its network states, thereby enabling it to switch among different attractors. In order to avoid that the network gets locked in a single one of these attractors, we add an adaptation property to neurons in CA3: If a neuron has fired continuously over 10 time steps, it cannot fire again during the subsequent 50 time steps.

We include an algorithmic description (Algorithm 1) in the Supplement for further implementation details.

For testing bottom-up unbinding with ambiguous cues (see Fig. 5), the experiment is based on binding 1 cue from 8 words, where the word length is *L* = 6000. Consequently, the input layer size is set at 48,000, the CA3 layer size at 24,000, and the memory layer size at 12,000. Synapses from the input to CA3 and from CA3 to the memory layer are subject to the standard BTSP rule with *f*_*q*_ = 0.0025 for the probability of a plateau potential in the postsnaptic neuron. Instead of a fixed threshold, a k-Winners-Take-All (k-WTA) mechanism for k = 60 is employed for determining which neurons in the CA3 layer fire (i.e., assume value 1). The threshold for neurons in the memory layer is fixed at 24, with no threshold reduction required during bottom-up unbinding since the input strength from the recurrent activity in CA3 is sufficient. We trained the model on a dataset of 1000 sentences and obtained the results presented in Fig. 5E and Fig. 5F. To construct the dataset containing 1000 items, with each group of four sharing the same cue word in the same position, we first randomly select 250 cue words for repetition. Then, each of these cue words is placed in the selected same position of the sentence. For each cue word, we generate 4 sentence completions by filling in the remaining 7 positions with randomly chosen words from the entire vocabulary of 1000 words. We measured the correctness of a network state by *max*(100% − *error*%, 0), where the error percentage was defined as before as ratio of the number of error bits and the number of expected 1’s. Since it is possible for this error measure that there are more error bits than the expected number of 1’s, we imposed a lower bound of 0 for the resulting correctness values. We evaluated bottom-up unbinding performance using datasets of 1000 and 4000 samples, with the correctness distribution shown in Fig 4G. We compared in Fig. 5E, F the network states that occurred in the course of the network dynamics during unbinding with those network states that resulted when each of the 4 possible sentence solutions were initially presented as network inputs during learning, without activating the recurrent connections in the CA3 module.

### 4.6 Details of iterated binding

#### 4.6.1 Details of hierarchical iterated binding (Fig. 6)

In hierarchical iterated binding, the encoding utilizes a hierarchical binary tree structure to organize the binding process, until a single top-level composed representation is achieved. For example, with four words, the process involves pairing and binding *A* and *B* to form ⟨*A, B* ⟩, and *C* and *D* to form ⟨*C, D*⟩ . These pairs are then bound together to form ⟨⟨*A, B*⟩, ⟨*C, D*⟩⟩, etc. This method can also be used for binding a list of words whose number is not a power of 2. In cases where a path contains only a single word, no binding operation occurs, and the word is directly propagated to the subsequent hierarchical level. We include an algorithmic description of hierarchical binding (Algorithm 2) in the Supplement S1. for further details.

We assume that a single network is used for binding and unbinding at each position in the iterated binding schemes, both for hierarchical binding and online binding. The feedforward weights of this network are trained in an online manner, for each input that is presented to the network. Symmetrically, feedback connections are adjusted together with the feedforwards weights at each of these learning steps. The best unbinding performance is likely to be achieved when it occurs right after the weight adjustments for this particular ensemble of input words. But our results in Fig. 6 C, D and Fig. 7 C, D show that unbinding still works very well after the weights of the network have been trained for many subsequent ensembles of input words. In other words, learnt performance for a single ensemble of input words is not ruined by subsequent learning processes for other ensembles of input words. The CAM for cleaning up the words that result from top-down unbinding at the input level was trained on the vocabulary of words that we employed.

In the experiments of both Fig. 6C and Fig. 7C, the word length is set at 12,000, with each word containing a fixed 60 one-bits (*f*_*p*_ = 0.005), forming a vocabulary of 1,000 words (*N*_*w*_ = 1000). Under the iterated binding model, the size of the memory layer is set equal to the word length, ensuring that the dimensions of the composed representation match those of the word, which allows for uniform processing of both word representations and composed representations. The dimension of the input layer is set to twice the word length to handle two representations simultaneously. The same model is reused at each binding step, with the size of the input layer set at double *L* (24,000) and the memory size equal to *L* (12,000), with *f*_*q*_ = 0.005 for BTSP maintained throughout. At the memory layer, during binding, the k-WTA strategy is used to select the 30 neurons with the highest activations to produce spikes (k=30 for k-WTA), ensuring a consistent number of spikes across all composed representations. During unbinding, activations after the backward pass are split evenly into two parts, each using k-Winners-Take-All (k-WTA) to achieve the expected number of one bits. Specifically, 30 neurons are selected for composed representations (k=30 for k-WTA), and 60 neurons for recalled words (k=60 for k-WTA). Note that similar performance can be achieved without WTA competition if the neuron thresholds are properly adjusted for the activation level at each binding and unbinding step. The CAM trained on the vocabulary is used after the unbinding process concludes. The input size for the CAM is 12,000, and the memory size is 24,000, where k-WTA selects 30 neurons in the memory layer for composed representations, and 60 neurons in the input layer for clean-up words. In both Fig. 6C and Fig. 7C, top-down unbinding results are depicted for varying ensemble lengths and numbers. The range of ensemble counts starts from 100 and extends to 6000, with an interval of 100 between each data point. The error on the z-axis represents the ratio of wrong bits to expected ones.

#### 4.6.2. Details of online iterated binding (Fig. 7)

In the online iterated binding (Fig. 7A), we start by combining two elements to form their composite (e.g., *A* and *B* to ⟨*A, B*⟩), and the composite ⟨*A, B*⟩ is then remapped back to the input layer, where it is combined with another element *C* to generate ⟨⟨*A, B*⟩, *C*⟩ . The process is repeated until the entire sentence is in its complete binding form. Given the number of operations performed by the network as depth, the word quantity contained in the compressed sentence is *K* = *depth* + 1. During the unbinding phase, we utilize the feedback weight that is pre-trained in the forward pass to achieve the mapping from composed representation to inputs recursively (Fig. 7B). We first start from the compressed form of the sentence to obtain the initial unbinding result (e.g., ⟨⟨⟨*A, B*⟩, *C*⟩, *D*⟩ to ⟨⟨*A, B*⟩, *C*⟩*′* and *D*^*′*^). Subsequently, ⟨⟨*A, B*⟩, *C*⟩*′* is re-introduced as an input in memory neurons, resulting in ⟨*A, B*⟩*′* and *C*^*′*^, and so forth, until we obtain the final unbinding results *A*^*′*^, *B*^*′*^, *C*^*′*^, *D*^*′*^.

## Acknowledgments

We thank Paxon Frady, Denis Kleyko, and Fred Sommer for helpful discussions. The research of WM was partially supported by the National Science Foundation of the USA (EFRI BRAID project 2318152) and the Austrian Science Fund (FWF) (10.55776/COE12). The research of Chengting Yu during his period at TU Graz was supported by the Feiying Program from Zhejiang University. The research of Aili Wang was supported by the National Natural Science Foundation of China under Grant No. 62304203. The authors gratefully acknowledge the Gauss Centre for Supercomputing e.V. (www.gauss-centre.eu) for funding this project by providing computing time on the GCS Supercomputer JUWELS[1] at Jülich Supercomputing Centre (JSC).

## Supplementary Information

### S1. Algorithms for BTSP-based binding

#### Algorithm 1

Combining binding in CA1 with association in CA3

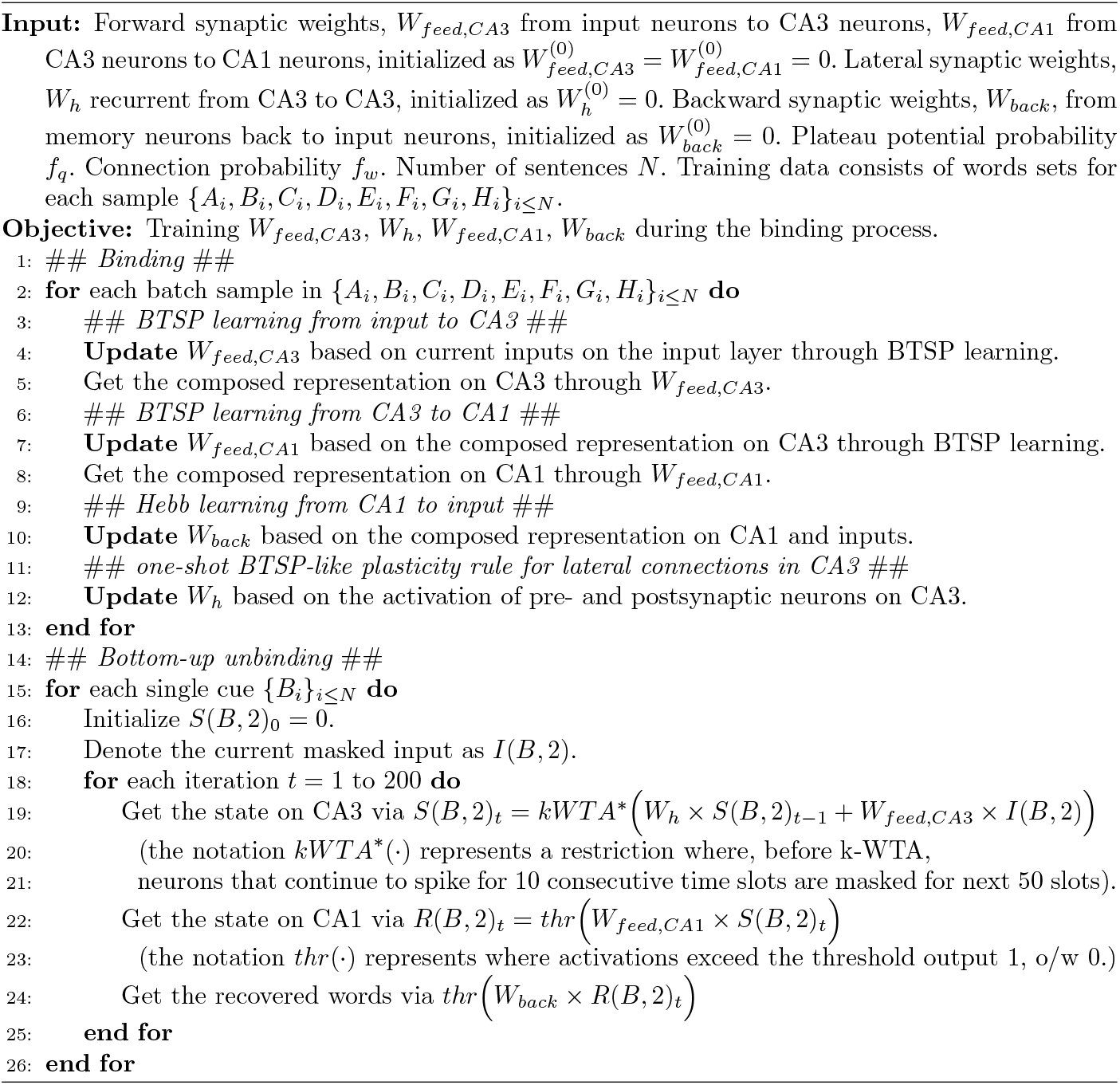

**Algorithm 1: Combining binding in CA1 with association in CA3**. We introduce CA3 as an intermediate layer to temporally decouple the superposition, allowing the network to sequentially oscillate among all valid patterns. Dense lateral connections and global k-Winners-Take-All (k-WTA) operations within CA3 enable attractors to form before propagating activation to the memory layer. During the binding phase, activity can pass through the CA3 layer without engaging the internal dynamics produced by its recurrent lateral connections. We apply one-shot synaptic plasticity to the weights *W*_*h*_ only once (Line 6), targeting the representation generated at the first step. We select the composed representations of CA3 (Line 4) and CA1 (Line 9) during the binding process as the states for subsequent comparisons between *S*(*B*, 2)_*t*_ and *R*(*B*, 2)_*t*_. The representation generated on Line 4, referred to in the caption of Fig. 5E, is recognized as the first state of the recurrent network module when a full sentence is presented on the input layer. The representation generated on Line 9 corresponds to the first state of the memory neurons when a full sentence is presented on the input layer. To ensure stability in the recurrent process, we employ k-Winners-Take-All (k-WTA), selecting the k neurons with the highest activation levels to fire spikes while the rest remain inactive, to control the number of spikes in CA3 (Line 17), where each activation consistently engages 60 neurons (k=60 for WTA).

#### Algorithm 2

Hierarchical iterated binding with eight words

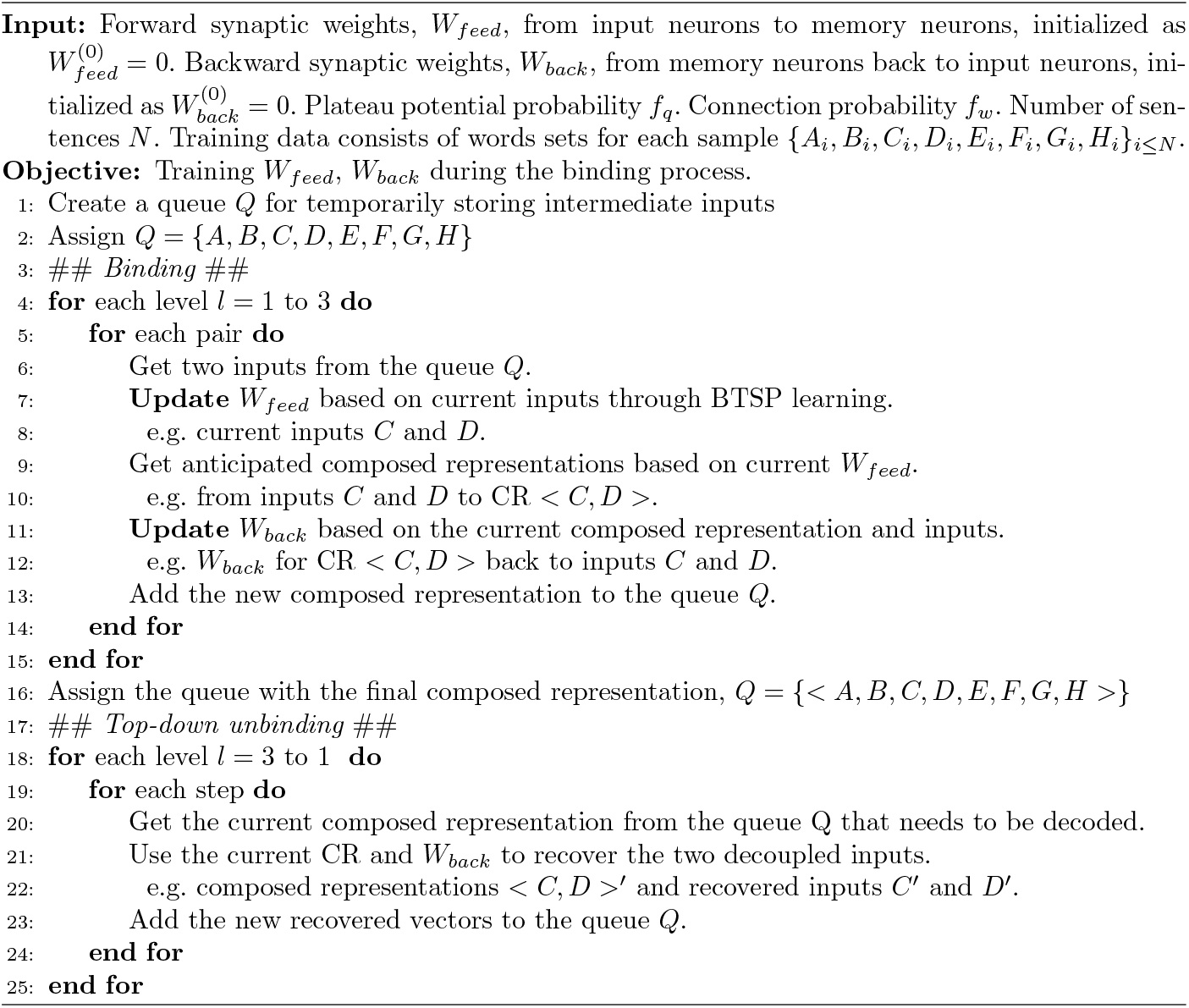

**Algorithm 2: Hierarchical iterated binding in the eight-words case**. Global weights *W*_*feed*_ and *W*_*back*_ are utilized throughout the entire binding process. For learning *W*_*feed*_ based on BTSP, the training protocol initiates at the first level with binding {*A, B*} into *< A, B >*, followed by binding {*C, D*} into *< C, D >* at the second step, and so forth. At the second level, the first step involves binding *< A, B >* and *< C, D >* into *<< A, B >, < C, D >>*. This process continues in the same manner until the final level, resulting in *<<< A, B >, < C, D >>, << E, F >, < G, H >>>*. The learning of *W*_*back*_ follows sequentially through the binding steps, mirroring the *W*_*feed*_ training. The training concludes at the final level, after which the network’s weights are finalized. At this point, the top-down unbinding process employs the network’s *W*_*back*_ to calculate the reversal from the final composed representation back to all intermediate composed representations and the input words.

### S2. Control experiments on sparsity

**Fig. S1:**
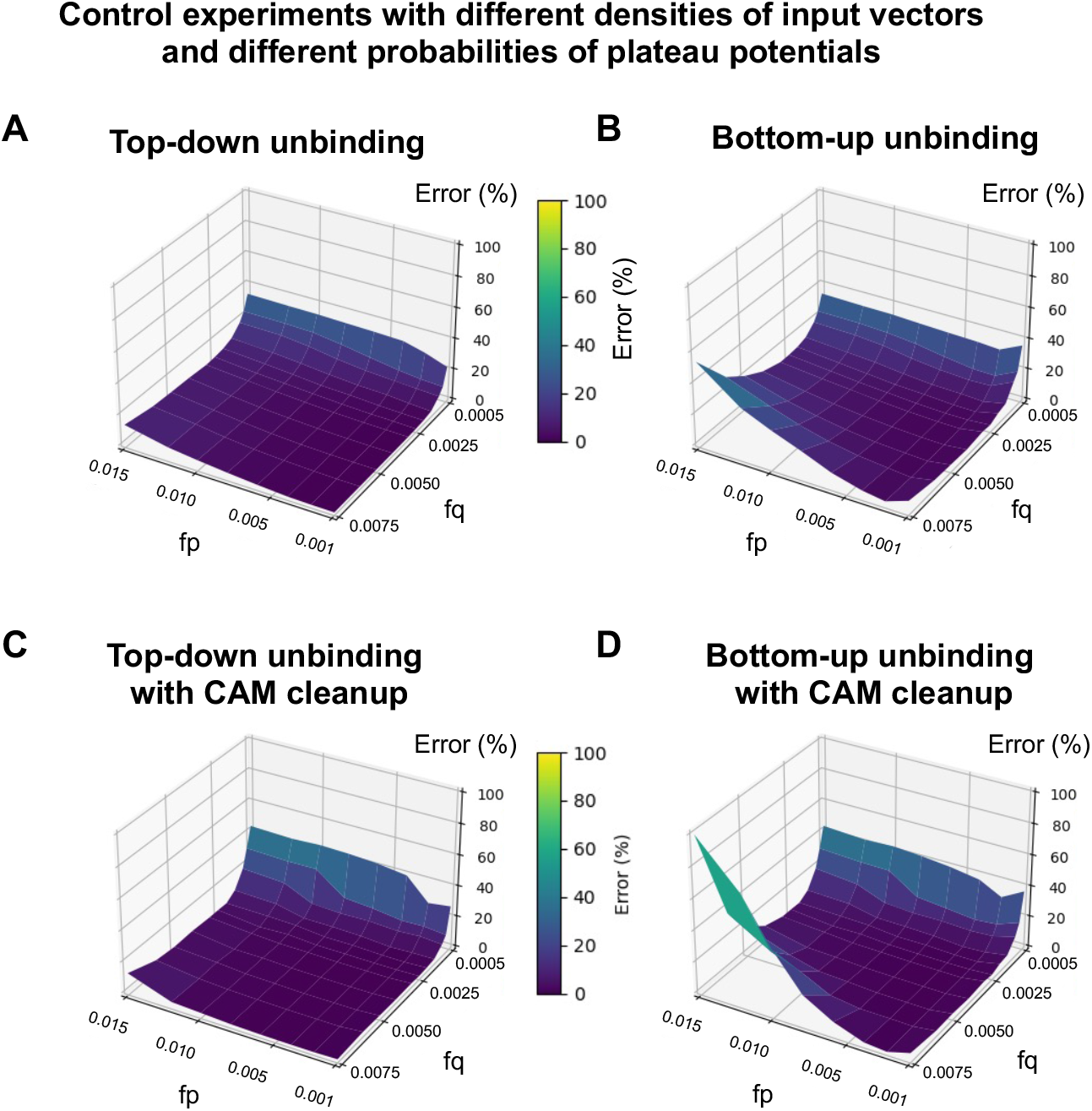
Control experiments on sparsity.

## Notes

### Competing Interest Statement

The authors have declared no competing interest.

### Summary of Updates

the new section 2.5 updated; the abstract, introduction, section 2.2, discussion, captions of Figures 9 and 10, and section 4.4 revised.

